# TimiGP: inferring inter-cell functional interactions and clinical values in the tumor immune microenvironment through gene pairs

**DOI:** 10.1101/2022.11.17.515465

**Authors:** Chenyang Li, Baoyi Zhang, Evelien Schaafsma, Alexandre Reuben, Jianjun Zhang, Chao Cheng

## Abstract

Determining how immune cells functionally interact in the tumor microenvironment and identifying their biological roles and clinical values are critical for understanding cancer progression and developing new therapeutic strategies. Here we introduce TimiGP, a computational method to infer inter-cell functional interaction networks and annotate the corresponding prognostic effect from bulk gene expression and survival statistics data. When applied to metastatic melanoma, TimiGP overcomes the prognostic bias caused by immune co-infiltration and identifies the prognostic value of immune cells consistent with their anti- or pro-tumor roles. It reveals the functional interaction network in which the interaction X→Y indicates a more positive impact of cell X than Y on survival. This network provides immunological insights to facilitate the development of prognostic models, as evidenced by our computational-friendly, biologically interpretable, independently validated models. By leveraging single-cell RNA-seq data for specific immune cell subsets, TimiGP has the flexibility to delineate the tumor microenvironment at different resolutions and is readily applicable to a wide range of cancer types.

## Main

Cancers are composed of not only tumor cells but also various non-cancerous cells that constitute a complex tumor microenvironment. Immune cells are one of the essential non-malignant tumor components that interact with each other to regulate cancer evolution^1, 2^. These cells, including cytotoxic cells such as CD8+ T cells^3^ and natural killer (NK) cells^4^, as well as immunosuppressive cells such as regulatory T cells (Treg)^5^ and myeloid-derived suppressor cells (MDSC)^6^, form a tangled tumor immune microenvironment (TIME). Recently, accumulating evidence has suggested that the TIME drastically impacts the clinical outcomes of cancer patients^7–10^. However, due to the immune cell crosstalk and co-infiltration, analyses of their impact on clinical outcomes have often led to conclusions contradictory to each other or opposite to their established anti- or pro-tumor roles^9^. Therefore, a comprehensive understanding of how the TIME influences the biological and clinical behaviors of cancers is needed to facilitate the development of novel therapeutic strategies.

To investigate the TIME, traditional immunophenotyping approaches, such as immunohistochemistry (IHC) and flow cytometry, are widely used in clinical practice. However, these methods are only able to analyze a small number of markers; therefore, only a few cell types can be assessed simultaneously. Single-cell RNA sequencing (scRNA-seq) has emerged as a powerful technique that enables transcriptomic profiling and cell subtyping at a higher resolution^11^. However, it is impractical in clinical practice due to its high cost and requirement of high sample quality. For example, formalin-fixed paraffin-embedded (FFPE) samples, the most commonly used clinical samples, are not amendable for scRNA-seq, not to mention that recovery can be cell type-specific and can distort expression profiles during tumor dissociation(ref)^12^. By contrast, RNA-seq is amendable for low-quality samples and can portray the transcriptome profile from bulk tissues, including various cell types. RNA-seq-based assays have been widely utilized for molecular profiling in clinical practice^13, 14^. Therefore, bulk transcriptomic profiling retains its advantage as an important approach to studying the TIME, due to its mature technologies, tissue availability, and low cost.

Over the last two decades, multiple marker expression- or deconvolution-based approaches (e.g., xCell^15^, CIBERSORTx^16^) have been proposed to dissect cellular heterogeneity in the TIME^15–25^. However, most of these methods are designed to estimate immune cell infiltration levels rather than immune interactions. Though a few methods enable the inference of cell-cell communication from bulk sequencing data (e.g., CCCExplorer^26^, ICELLNET^27^), these require knowledge of ligand-receptor interactions and hence limit their application only to known intercellular signaling communications^28^. Due to our rudimentary knowledge of the TIME, these methods often face challenges in defining the roles of various immune cell types during cancer evolution, complex inter-cell interactions, and their association with clinical outcomes. Therefore, computational methods that comprehensively dissect TIME functional networks and assess their clinical values remain an unmet need.

To fill this void, we developed TimiGP (<u>T</u>umor <u>I</u>mmune <u>M</u>icroenvironment <u>I</u>llustration based on <u>G</u>ene <u>P</u>airing), a computational method to investigate the TIME by inferring inter-cell “functional” interactions and the prognostic value of immune cells. Our approach combines survival statistics with bulk transcriptomic profiles to construct a “functional” immune interaction network. The network reveals two “functions” of immune cells: 1) biological function, the anti- or pro-tumor capacity; 2) clinical function, the prognostic values of immune cells. In this study, we applied TimiGP to metastatic melanoma and demonstrated how TimiGP resolves the prognostic bias (i.e., the estimation of the clinical value of immune cells contradictory to their biological function) caused by immune co-infiltration. Referring to the functional interaction network, we also built a prognostic model to denote the clinical utility of TimiGP, which exhibited high interpretability and accuracy across multiple independent datasets. To extend the application of TimiGP, we integrated single-cell RNA-seq (scRNA) results and enabled the method to analyze different scales of inter-cell functional interactions, such as in the entire tumor microenvironment or within T-cell subpopulations. Finally, we applied TimiGP to 23 solid tumor types and revealed the clinical value of immune cells in pan-cancer. This work will improve the understanding of the TIME with implications for novel target identification and prognostic model construction, eventually facilitating personalized therapies.

## Result

### Gene pair analysis disentangles expression correlations caused by immune cell co-infiltration

Due to inter-cell crosstalk, different immune cells are often closely regulated and hence co-infiltrate the tumor microenvironment^29^. To demonstrate this, we applied eight transcriptome-based cell-type quantification methods^15–25^ to metastatic melanoma, which is known to be a highly immune infiltrated cancer type (immune “hot”)^30,31^. As expected, estimated cell abundances were strongly positively correlated to each other (Figure S1A). Although co-occurrence patterns can be used to identify multicellular communities (e.g., EcoTyper), they may also cause “prognostic bias” when the clinical significance of immune cells is inappropriately estimated. We examined the association between prognosis and inferred immune cell infiltration (Table S1A). As shown in Figure 1A, 44 cell types were significantly associated with prognosis (P-Value < 0.05), with the majority (93.2%) associated with a favorable prognosis (Hazard Ratio < 1), including many well-known pro-tumor cell types including tumor-associated neutrophils^32^ and plasmacytoid dendritic cells^33^. This counterintuitive association between the high infiltration of pro-tumor cells with superior survival is an instance of prognostic bias. Similarly, at the gene expression level, most prognostic immune marker genes (IMGs) (90.5%), including many negative immune regulators (e.g., *PD-L1, LAG3, TIGIT, IDO1*)^34, 35^, were associated with improved survival (Figure 1B, Table S1B), likely also due to a high level of immune co-infiltration as evidenced by the predominant positive correlation between their IMGs (Figure 1C, Figure S1B). Taken together, these results highlight the need for new methods to reduce the potential influence of immune co-infiltration, in order to reveal the true clinical and biological roles of immune cells.

**Figure 1.**
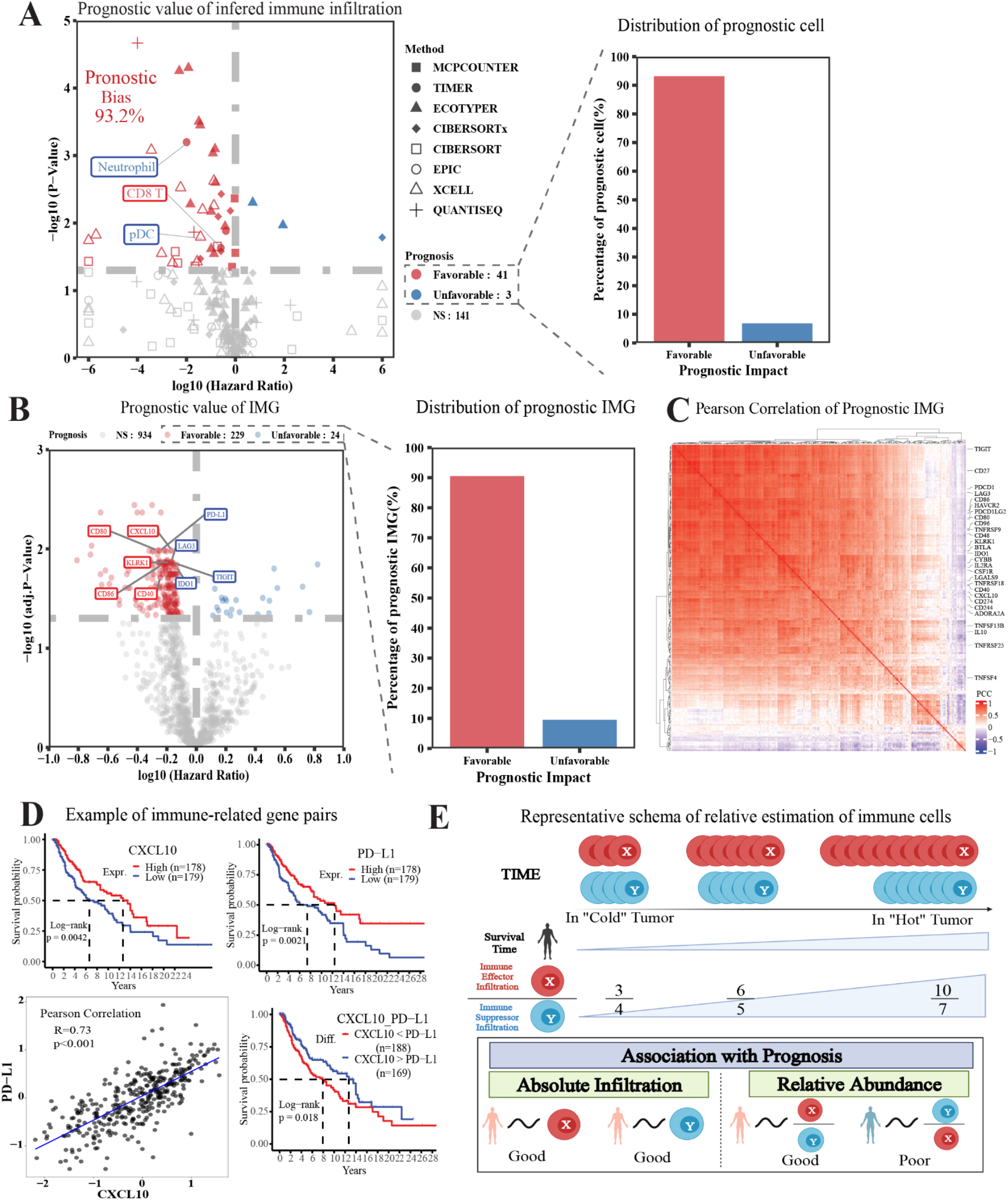
Gene Pairs was a potential solution of prognostic bias caused by immune cell co-infiltration. (A) Volcano plot showing log-transformed hazard ratio(x-axis) and P values(y-axis) in the Cox regression model that fits infiltration of each cell type as a continuous variable. The cell type is defined, and the infiltration level is estimated by eight methods: MCP-counter^17^, TIMER^18–20^, EcoTyper^21^, CIBERSORTx^16^, CIBERSORT^22^, EPIC^24^, xCELL^15^, and quanTIseq^23^. The percentage of cell types associated with favorable(red) or unfavorable(blue) prognosis is stressed in the bar graphs. (B) Volcano plot showing log-transformed hazard ratio(x-axis) and BH-adjusted P values(y-axis) in the Cox regression model that fits the expression of each immune marker gene(IMG) as a continuous variable. The text labels well-known immune stimulators/cytotoxic markers(red) and inhibitors/checkpoints(blue). The percentage of IMGs associated with favorable(red) or unfavorable(blue) prognosis is stressed in the bar graphs. (C) Heatmap of Pearson correlation coefficients(PCC) of prognostic IMGs’ expression. (D) Example of PD-L1 and CXCL10 showing that the high correlation between the two genes’ expression leads to prognostic bias, and their gene pair can solve the bias. Upper panel: Kaplan-Meier(KM) curve showing the overall survival of patients with high(red) or low(blue) expression of CXCL10 (left) or PD-L1(right) using the median as a cutoff; Lower panel: Scatter plot showing the Pearson correlation between the two genes(left) and KM curve showing the survival impact of CXCL10-to-PD-L1 ratio (right). The P-value shown in the KM plot is calculated by the log-rank test by comparing the survival of two groups. (E) Representative schema showing how the correlated infiltration affects the prognostic evaluation of cells and indicating the relative estimation(e.g., relative abundance) is a better measurement than absolute infiltration. All analysis has been performed with the RNA-seq of metastatic melanoma in the Cancer Genome Atlas(TCGA_SKCM06)^39^. See also Figure S1 – S2.

Immune-related gene pairs have been reported as predictive biomarkers in previous studies^36, 37^. Theoretically, the co-expression of IMGs can be de-correlated by focusing on the relationship between pairs, thereby reducing the prognostic bias (Figure 1D, Figure S2). For example, in Figure 1D, both *CXCL10* and *PD-L1* were associated with longer survival, although they have opposing functions - promoting (CXCL10) versus suppressing (*PD-L1*) anti-tumor immunity, respectively. The positive prognostic value of PD-L1 is likely caused by its positive correlation with genes promoting anti-tumor immunity (e.g., *CXCL10,* which increases T cell infiltration and IFN-γ production and causes *PD-L1* expression^38^). Interestingly, a higher *PD-L1*-to-*CXCL10* ratio was associated with significantly shorter survival, suggesting that the relative expression level of immune genes may better reflect the biological/clinical behaviors of cancers. Inspired by such gene pairs, we considered estimating the relative abundance at the cellular level to reduce the influence of co-infiltration and therefore inferring the prognostic value of immune cells more accurately. As shown in Figure 1E, although the absolute infiltrations of immune effectors and suppressors are positively correlated, their relative abundance enables us to capture subtle differences between them and reveal the prognostic values of these cells in line with their biological functions. As an extension of this idea, we developed a novel method called TimiGP, Tumor Immune Microenvironment Illustration based on Gene Pairing.

### The TimiGP framework

TimiGP is designed to illustrate the TIME by inferring functional interaction networks and annotating clinical values of infiltrating immune cells from bulk tissue transcriptomes. To achieve this, it requires IMG expression profiles and patient survival statistics as input. TimiGP consists of five major steps: 1) performing pairwise comparisons between IMGs based on expression profiles, 2) selecting prognostic IMG pairs (IMGP) by survival analysis, 3) generating cell interaction annotations from cell-type markers, 4) determining functional interactions with enrichment analysis, and 5) annotating the clinical function of immune cells through network analysis (Figure 2; Methods).

**Figure 2.**
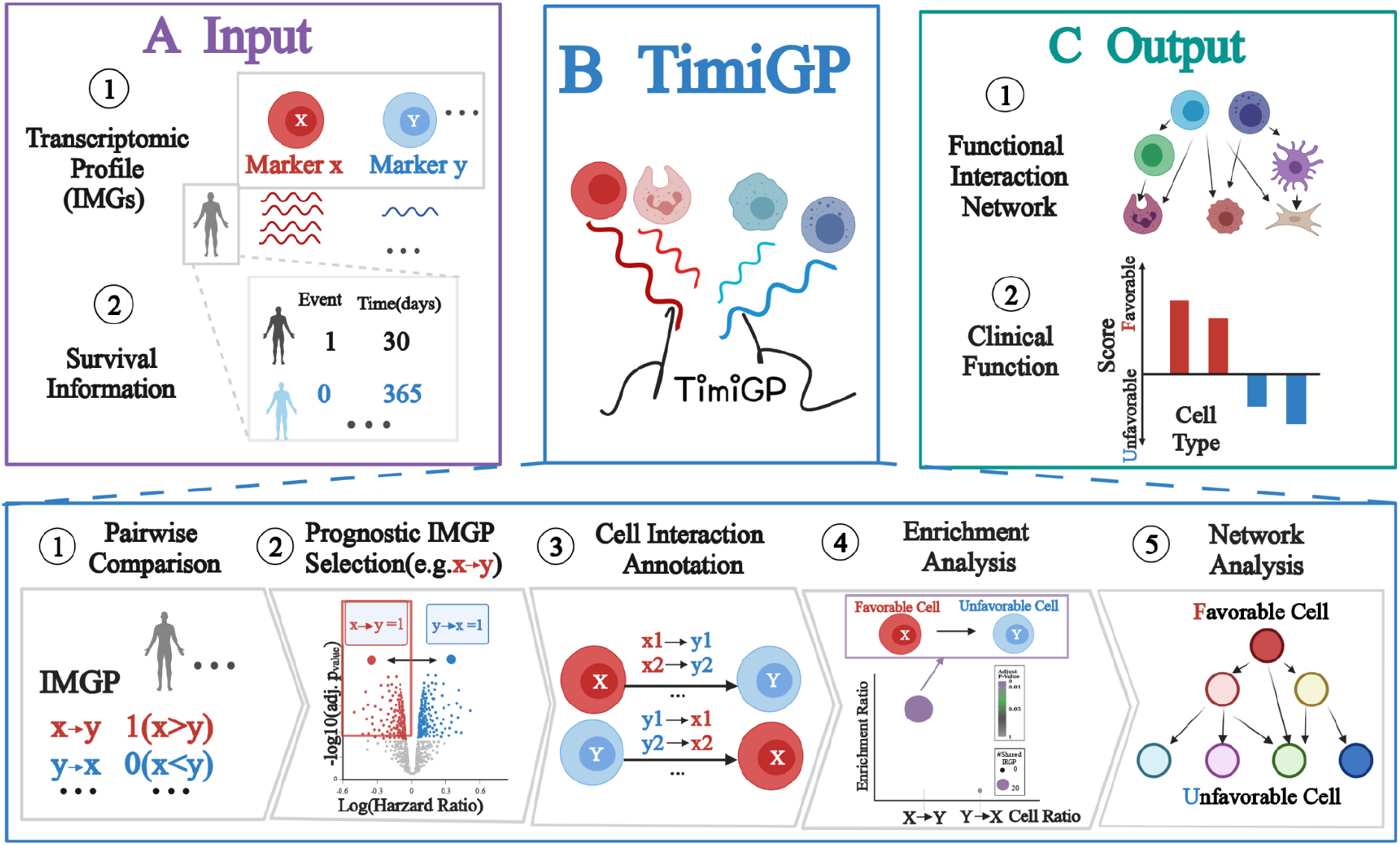
TimiGP framework. Schematic depicting the TimiGP framework. TimiGP required two inputs: 1) transcriptomic profile of immune marker genes(IMGs); 2) survival statistics including event(e.g., death, recurrence) and time-to-event of the same cohort. TimiGP will infer inter-cell functional interactions and estimate prognostic values of immune cells through five steps: 1) performing pairwise comparisons between IMGs based on expression profiles, 2) selecting prognostic IMG pairs by survival analysis, 3) generating cell interaction annotations from cell-type markers, 4) determining functional interactions with enrichment analysis, and 5) annotating the clinical function of immune cells through network analysis.

In detail, TimiGP first captures the logical relation between any two IMGs according to bulk tissue transcriptomes (Step 1). For example, if the expression of gene x is higher than y, the marker pair score (MPS) is 1; otherwise, the MPS is 0. In this way, it evaluates the relative abundance between IMGPs in an individualized manner, which ensures that subsequent steps are not affected by immune co-infiltration. Second, TimiGP employs cox regression to evaluate associations between IMGPs and prognosis (Step 2). It selects prognostic IMGPs whose MPS of 1 is significantly associated with a favorable prognosis (Hazard Ratio < 1; Benjamini-Hochberg(BH) adjusted P-value < 0.05). For example, a prognostic IMGP, x →y, indicates that a higher expression of marker x over y positively impacts survival. Third, TimiGP generates a cell interaction annotation to evaluate the relative strength of cell function from gene pairs (Step 3). Assuming the TIME is a balance between pro- and anti-tumor immunity, which of these two prevails determines the capacity of the immune system to promote versus inhibit cancer progression and subsequently impacts prognosis accordingly. We simplified the complex TIME balance to the relative function between paired immune cells, whose functions are represented by their marker genes. For example, the cell X → cell Y interaction is determined by combining cell X markers with cell Y markers (e.g., x1→y1, x2→y2). If more cell X-to-Y markers than cell Y-to-X markers are prognostic pairs, it suggests a more powerful cell X over cell Y function associated with favorable clinical outcomes. Accordingly, we defined cell X → cell Y as a “functional” interaction. The “function” describes the aspects of immune cells: 1) Cell function: the relative effectiveness between cells’ ability to inhibit or promote tumor progression, which can be further explained by the relative abundance between cell infiltration or/and the marker expression per cell. 2) Clinical function: the relative favorability of immune cells in prognosis. To statistically determine the functional interaction, TimiGP then performs the enrichment analysis (Step 4). Suppose prognostic IMGPs significantly share more IMGPs with cell interaction X→Y annotation than is expected by chance. In that case, the cell X→Y interaction is selected as a functional interaction (BH adjusted P-value < 0.05), which denotes that the more effective X function than Y positively impacts survival. Correspondingly, cell X tends to be an anti-tumor cell associated with a favorable prognosis, while cell Y is likely to be a pro-tumor cell associated with an inferior outcome. Based on enrichment results, TimiGP integrates all functional interactions to construct a directed network and scores the clinical roles of immune cells according to the network degrees (Step 5).

### TimiGP illustrates clinically relevant TIME in metastatic melanoma

To demonstrate the capabilities of TimiGP, we applied our method to metastatic melanoma cohorts, where, as mentioned, immune co-infiltration causes prognostic bias. We started with IMGs of 22 cell types identified in immunoscore, of which the performance has been extensively validated and supported^40^. In addition, we kept markers of two special types: the cytotoxic cells denoted by common cytotoxic markers of, for example, anti-tumor CD8 T cells and NK cells, as well as the tumor cells, as evidence of immune evasion. Both unique cell types were set as positive and negative controls, respectively.

Among the 462 possible pairwise cell interactions, TimiGP identified 54 as functional interactions (Table S2). Cytotoxic cells →Mast cells, Cytotoxic cells →Neutrophil, Cytotoxic cells →Tumor cells, and Cytotoxic cells →Immature dendritic cells (iDC) were the first four functional interactions ranked by BH-adjusted P-value (Figure 3A), suggesting that the anti-tumor function (from cytotoxic cells) stronger than the pro-tumor function of the second cell types may have contributed to a favorable prognosis. Significantly, the interaction from the positive control (cytotoxic cells) to the negative control (tumor cells) is circled out, which attests to the validity of the method. Similarly, the Type 1 helper T cell (Th1) also stood out as a favorable cell among the top 10 functional interactions.

**Figure 3.**
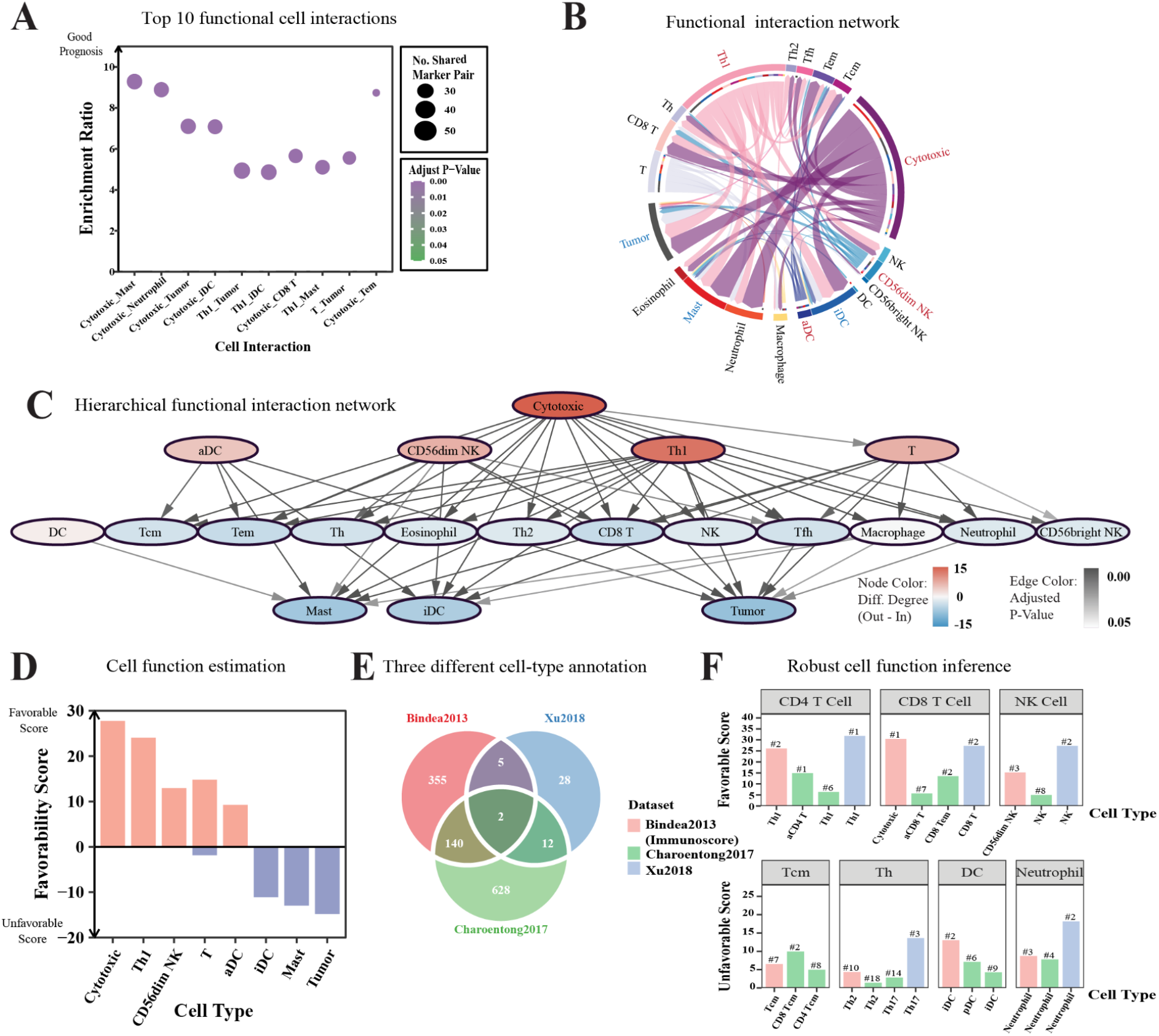
TimiGP robustly infers immune cell interactions and their favorability on prognosis. (A - D) Application of TimiGP to metastatic melanoma based on modified immunoscore markers(Bindea2013)^40^. (A) Dot plot of the top 10 functional interactions ranked by BH-adjusted p-value. (B) Chord diagram of all functional interactions. The arrow represents functional interactions from favorable cell type(color of the outer ring and arrow) to unfavorable cell type(color of the inner ring). The wider the arrow is, the smaller the adjusted p-value is. (C) A hierarchical functional interaction network based on degrees. The node represents the cell type, and its color shows the difference between out-degree and in-degree. The edge represents functional interaction, and its transparency shows the BH-adjusted p-value. (D) Bar plot of the favorability score to evaluate each cell type’s favorable(orange) or unfavorable(blue) role in anti-tumor immunity and prognosis. (E) Venn diagram showing the numbers and overlap of IMGs in 3 immune cell marker annotations. (F) Bar plot of the favorable score(upper) and unfavorable score(lower) of similar cell types defined in 3 different marker annotations: Bindea2013^40^(red), Charoentong2017^41^ (green), Xu2018^42^(blue). The text label shows the rank of the cell favorability in each annotation set. The non-immune cell type, tumor cell, has been removed in Bindea2013 annotation before TimiGP analysis. All analysis has been performed with the RNA-seq of metastatic melanoma in the Cancer Genome Atlas(TCGA_SKCM06). Abbreviation of cell types: T, T cell; Th, Helper T cell; Th1, Type 1 Th; Th2, Type 2 Th; Tfh, Follicular Th; Tcm, Central memory T cell; Tem, Effector memory T cell; aCD4 T, Activated CD4 T cell; aCD8 T, Activated CD8 T cell; Cytotoxic, Cytotoxic cell(common cytotoxic markers of anti-tumor CD8 T cells, Tγδ, and NK cells)^40^; NK, Natural killer cell; DC, Dendritic cell; iDC, Immature DC; aDC, Activated DC; pDC, Plasmacytoid DC; Mast, Mast cell; Tumor, Tumor cell. See also Figure S3.

Using the 54 functional interactions, we constructed the network as one of the major TimiGP outputs. As shown in Figure 3B, activated dendritic cells (aDC), Th1, cytotoxic cells, and CD56dim NK cells were identified as favorable cell types compared to all other cell types, with mast cells, iDC, and tumor cells (negative control) as unfavorable cell types for prognosis, which was consistent with the known functions of these cells^3, 43–46^. Taken together, these results indicate that TimiGP can potentially overcome the limitation of immune co-infiltration and identify the real biological and clinical values of each immune cell subtype.

To quantify the prognostic value of these cells, we further analyzed the properties of the functional interaction network. This directed network demonstrates a hierarchical structure determined by the degree - the number of edges of one given node connected to other nodes (Figure 3C). The top layer (out-degree only) is composed of the favorable cells mentioned above, together with the positive control (cytotoxic cells), while the bottom layer (in-degree only) contains unfavorable cell types, as well as the negative control (tumor cells). Though the middle layer is more complicated as a mixture, this degree-dependent hierarchical network reveals the corresponding functional roles of various cell types in cancer biology and prognosis.

Considering that the out-degree and in-degree indicate one cell type to be more favorable or unfavorable than other types, we scored the prognostic value of each cell type by counting degrees. Since most immune cells have been reported to have dual roles in cancer evolution, we evaluated them from both sides: 1) “favorable scores” represent the anti-tumor function and positive prognostic value; 2) “unfavorable scores” represent the pro-tumor function and negative prognostic value. This “favorability score” is another primary output of TimiGP. As shown in Figure D, the result is consistent with network analysis and previous studies^3, 43–46^. The positive control (cytotoxic cells) demonstrated the highest favorable score, while the negative control (tumor cells) showed the highest unfavorable score. Altogether, these results confirm the validity of TimiGP for dissecting the TIME based on gene pairing rationale and identifying the reasonable prognostic values of immune cells by constructing the inter-cell functional interaction network.

We next sought to evaluate the robustness of TimiGP. Since the method relies on the quality of IMG markers, robustness here describes whether it remains reliable when using different cell-type annotations (Figure 3E). Compared to the cell-type markers used in the development, signatures generated by Charoentong, P. et al.^41^ include more mutually exclusive cell subtype markers, while modified IMGs are based on Xu. L et al.^42^, with only 47 genes used to represent 17 cell types. Though the cell classifications were distinct, TimiGP still identified CD8 T cells →Neutrophils (Figure S3A) and Th1 →Neutrophils (Figure S3B) from three analyses, which indicates the negative prognostic value of neutrophil-to-lymphocyte ratios, in line with recent studies^47–51^. Furthermore, consistently estimated with three analyses, activated T cells and NK cells contributed to the favorable prognosis, whereas neutrophils were relatively unfavorable (Figure 3F).

Moreover, we also used IMGs from CIBERSORT LM22^22^ for TimiGP to analyze the same cohort. Compared to CIBERSORT^22^ and CIBERSORTx^16^ (Figure 1C), TimiGP was able to identify the association of immune cells with clinical outcomes more consistent with their biological functions^2, 9, 44^. (Figure S3C, D). Collectively, TimiGP demonstrated stable performance when using different cell-type annotations to infer immune cell interactions and functions associated with prognosis.

### TimiGP facilitates the development of IMGP-based prognostic models with immunological insights

Immune-related gene pairs have been reported as emerging biomarkers to predict clinical outcomes^36, 37, 52^. Traditional methods select signatures from prognostic marker pairs, while TimiGP yields the enriched IMGPs that not only provide clinical insights but also reflect the functional cell interactions (Figure 4A), which is supposed to facilitate the model construction. Therefore, we next investigated the potential of TimiGP in the development of gene expression-based prognostic models.

**Figure 4.**
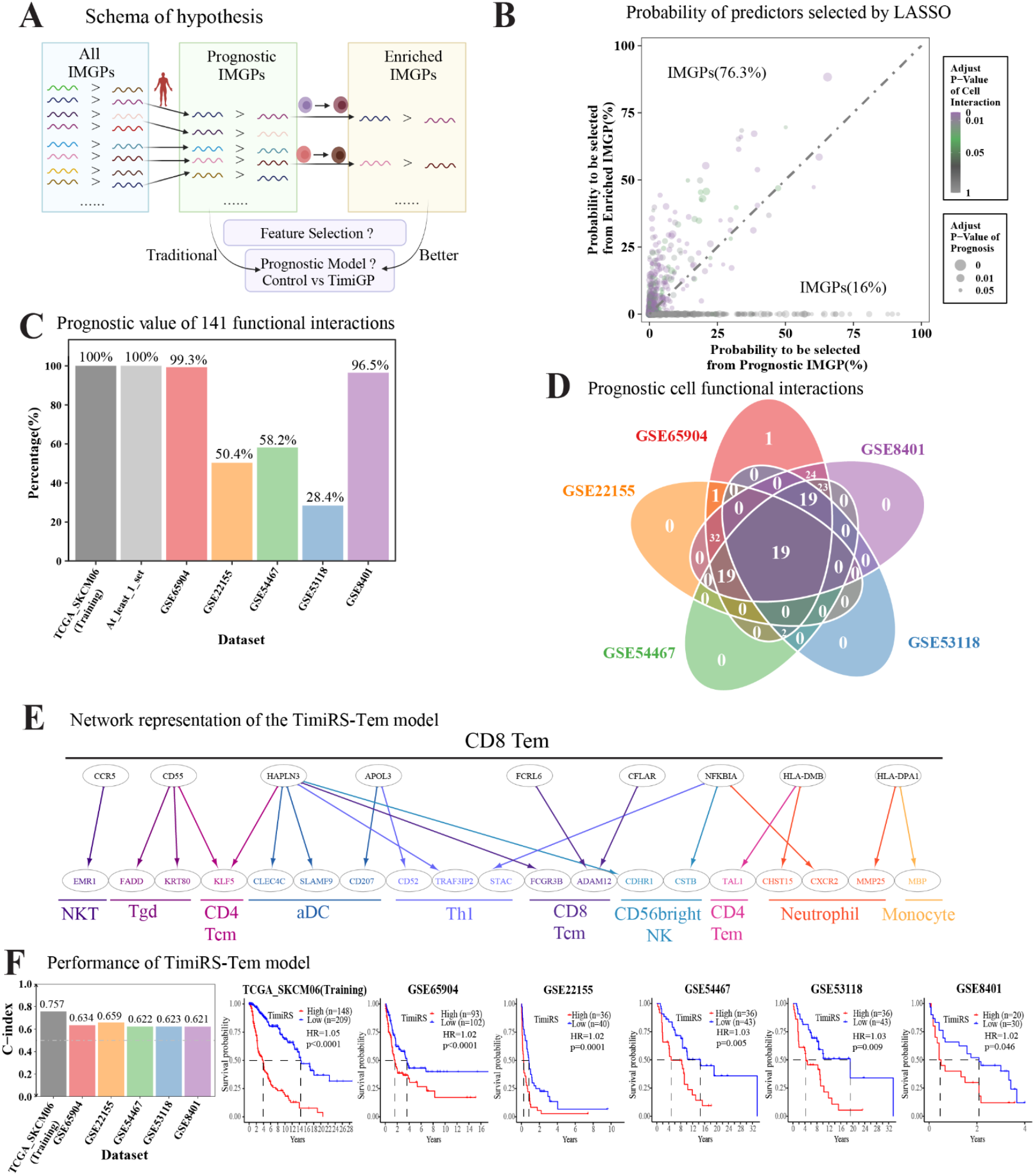
TimiGPS assists feature selection and IMGP-based prognostic model development. (A) Schema showing the rationale that TimiGP benefits feature selection with immunological insights and hypothesis that it facilitates the development of prognostic models. (B) Scatter plot of the probability of IMGPs picked from prognostic(x-axis) and/or enriched candidates(y-axis) by LASSO. The dot size shows the BH-adjusted P-value calculated by survival analysis, and the color shows the BH-adjusted P-value of potential cell interactions annotated by this IMGP according to enrichment analysis. The text label shows the percentage of both selected IMGPs in this area of 544 candidates belonging to prognostic and enriched IMGPs simultaneously. (C) Percentage of functional interaction-based TimiRS models that were statistically significant prognostic in training(TCGA_SKCM06) and five independent validation datasets (GSE65904^54^, GSE22155^55^, GSE54467^56^, GSE53118^57^, GSE8401^58^). “At_least_1_set” is the percentage of these models validated by at least one validation set. The prognostic value is evaluated by the univariate cox regression model that fits the TimiRS as a continuous variable, and the significance is p < 0.05. (D) Venn diagram showing the numbers and overlap of functional interaction-based TimiRS models validated by five independent validation sets. (E) A network representation of 21 pairwise features in the TimiRS-Tem model. Nodes represent immune marker genes. The color of the node label represents different immune cells. Edges describe pairwise relations. The edge from node x to node y denotes that a high x-to-y expression ratio is associated with better prognosis in patients whose color is the same as the color of the cell types at the lower layer. (F) Bar plot of C-index(left) and Kaplan-Meier curves(right) showing the performance of the TimiRS-Tem model. The Hazard Ratio and the corresponding P-value are calculated by the univariate cox regression model that fits the TimiRS as a continuous variable. Abbreviation of cell types in Charoentong2017 annotation^41^: Tem, Effector memory T cell; Th1, Type 1 Helper T cell; Tgd, Gamma delta T cell(Tγδ); Tcm; Central memory T cells; aDC, Activated dendritic cell; NK, Natural killer cell; NKT, Natural killer T cell. See also Figure S4.

We compared the probability that an IMGP was picked from enriched pairs with prognostic pairs through LASSO Cox regression^53^ (Methods). This time, we still utilized the same metastatic melanoma cohort but with the cell-type markers summarized by Charoentong, P. et al.^41^, which are mutually exclusive with more immune cell subtypes. As a result, there were 544 candidates taken from both of the input pools (Figure 4B, Table S3). Among these gene pairs, 76 % of IMGPs were more likely to be selected from enriched pairs than prognostic pairs to construct the model, suggesting that TimiGP benefits feature selection by filtering the correlated but noisy features.

In addition, the majority of model candidates denote high-confidence functional interactions between immune cells (Figure 4B, Table S3). This suggests that these functional interactions are potential signatures for prognostic models. To examine this, we designed TimiRS, the risk score calculated by the percentage of IMGP signatures with MPS of 0, and inspected the prognostic value of each functional interaction in one training set and five independent validation sets^54–58^ (Methods). The analysis revealed all functional interactions significantly associated with prognosis in at least one validation set (Figure 4C, Table S4), and 19 functional interactions were able to stratify survival across all five independent datasets (Figure 4D). These results suggest that TimiGP uncovers functional interactions of immune cells whose representative IMGPs show potential for both biological research and clinical applications.

Considering that single functional interactions are insufficient to portray the TIME, we further integrated several functional interactions and developed a new prognostic model. We noticed that the top functional and prognostic interactions were related to Effector memory CD8 T cells (CD8 Tem)-to-other cells, which supports the favorable role of CD8 Tem in anti-tumor immunity (Figure S4A). This is consistent with the characteristics of CD8 Tem, which undergo clonal expansion and exert anti-tumor function in the context of persistent antigen exposure^59, 60^. Therefore, we selected the top 21 IMGPs ranked by the LASSO selection probability to represent CD8 Tem → Other cells interactions and constructed the TimiRS-Tem model (Figure 4E, Table S5, Methods). As a control, 21 IMGP signatures were selected from prognostic pairs in the same way (Table S5). Compared to the control model (Figure S4B, C), TimiRS-Tem achieved a higher ability of patient stratification and concordance index (0.621-0.659) in five independent validation sets^54–58^ (Figure 4F). Its performance was consistently better than the control model when evaluated by the time-dependent receiver operating characteristic (ROC) curve, specifically AUC (the area under the curve) (Figure S4D, E). Taken together, these results suggest that TimiGP may facilitate the development of prognostic models by considering biological motivation and immunological dependency.

### TimiGP elucidates the tumor microenvironment at different scales

Over the past decade, single-cell RNA sequencing (scRNA-seq) has generated a large amount of cell-type annotations of different tissues at a granular level^61^. scRNA-seq-derived cell-type signatures have obvious advantages over bulk RNA-seq for their high specificity and resolution^62^. Therefore, we next sought to evaluate whether TimiGP was capable of dissecting the cell-cell interactions and functions at different scales by leveraging scRNA-seq-derived markers (Figure 5A).

**Figure 5:**
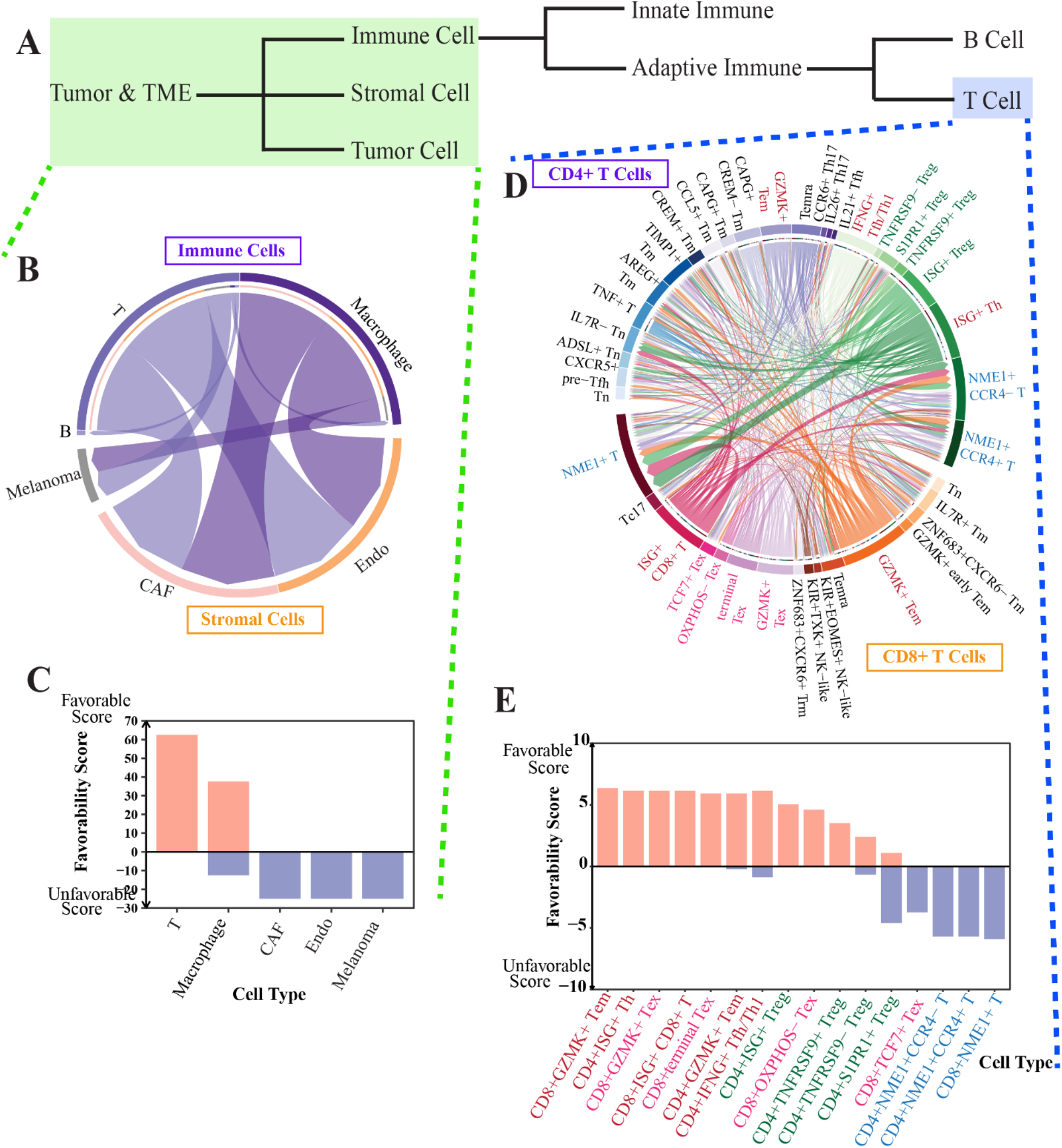
TimiGP can adjust the study resolution by utilizing cell-type markers from scRNA-seq. (A) A hierarchical schema showing tumor and microenvironment components. TimiGP was applied to metastatic melanoma(TCGA_SKCM06) to zoom out (green) and zoom in (blue) the tumor microenvironment. (B, C) Chord diagram of functional inter-cell interaction network(B) and the favorability score of selected cell types(C) to illustrate the entire tumor microenvironment using the markers summarized by scRNA-seq analysis on metastatic melanoma. (D, E) Chord diagram of functional inter-cell interaction network(D) and the favorability score of selected cell types(E) to illustrate the zoom-in T cell subpopulations using the markers summarized by scRNA-seq of pan-cancer T cell analysis. The color of cell types shows the top favorable(red) or unfavorable(blue) cell types, heterogeneous Tex(pink) and Treg(green). Abbreviation of cell types: CAF, Cancer-associated fibroblast; Endo, Endothelial cells; B, B cell; T, T cell; Tn, Naïve T cell; Th, Helper T cell; Th1, Typer Th; Th17, Type 17 Th; Tfh, Follicular Th; Treg, Regulatory T cell; Tex, Exhausted T cell; Tc17, Type 17 T cell; Tm, Memory T cell; Tem, Effector memory T cell; Trm, Tissue-resident memory T cell); Temra, Terminally differentiated effector memory or effector T cell.

We first applied TimiGP to dissect the entire tumor microenvironment of metastatic melanoma. In addition to immune cells, stromal cells are another critical component that shapes the microenvironment^63^. In 2016, Tirosh et al. applied scRNA-seq to 19 patients with metastatic melanoma and profiled tumor, immune, endothelial cells (Endo), and cancer-associated fibroblasts (CAFs)^64^. Using cell type markers identified in that study, we profiled the functional interactions between cancer cells and non-cancerous cells in the tumor microenvironment (Figure 5B). Compared to immune cells, cancer cells and stromal cells were associated with poor prognosis, consistent with the original scRNA-seq study^64^ (Figure 5C). This indicates that TimiGP can be utilized to study cell-cell interactions and functions which extend beyond cancer and immune cells.

Next, we tested the capability of TimiGP for high-resolution analysis. Using the T cell subtype markers identified from scRNA-seq analysis^65^, we repeated TimiGP in the same metastatic melanoma cohorts. Consequently, *GZMK+* effector memory CD8 and CD4 T cells contributed to a favorable prognosis, consistent with the analysis described above (Figure 5B, C). In addition, *IFNG+* follicular helper/Type 1 helper dual-functional T cells (Tfh/Th1) and ISG+ T cells, identified as potential tumor-reactive types in the original scRNA-seq analysis^65^, were also identified as anti-tumor cell types more strongly associated with a favorable prognosis than other cells. Our results also highlighted the heterogeneity of exhausted and regulatory T cells(Tex and Treg) from the perspective of clinical values. For example, the scRNA-seq study revealed that the *CD8+ TCF7+* Tex cell population had lower expansion and proliferation indices and a higher expression of exhausted and inhibitory markers, compared to *CD8+ GZMK7+* Tex^65, 66^. TimiGP demonstrated the heterogeneity of CD8+ Tex subtypes based on functional interactions and clinical values, which further supported that the *CD8+ GZMK7+* Tex cell was less favorable in prognosis than *CD8+ TCF7*+ Tex. In summary, TimiGP is capable of delineating the functional interaction network within subpopulations and identifying their potential functions at a more granular level using scRNA-seq.

Taken together, these results indicate that TimiGP is flexible in analyzing detailed cell subpopulations or broader tumor microenvironments, and TimiGP makes it possible to depict the TIME with high-resolution using low-resolution bulk sequencing data by leveraging high-resolution cell subtyping markers derived from scRNA-seq.

### TimiGP shows high potential in pan-cancer analysis

Finally, we applied TimiGP to 23 solid cancer types to showcase the generalization of TimiGP in studying the TIME (Table S6). As before, we set cytotoxic cells as the positive control and the tumor cells as the negative control. Although functional interactions vary across those cancer types, the impact of Cytotoxic cells →Other cells, or Other cells →Tumor cells interactions tends to remain consistent (Figure S5A, B). Furthermore, the prognostic value of some immune cells demonstrated similar patterns across most cancer types(Figure 6). For example, anti-tumor immune cells, such as B cells and cytotoxic cells, were more favorable regarding prognosis than tumor cells and pro-tumor immune cells, such as Neutrophils and Type 2 T helper cells across many cancer types.

**Figure 6:**
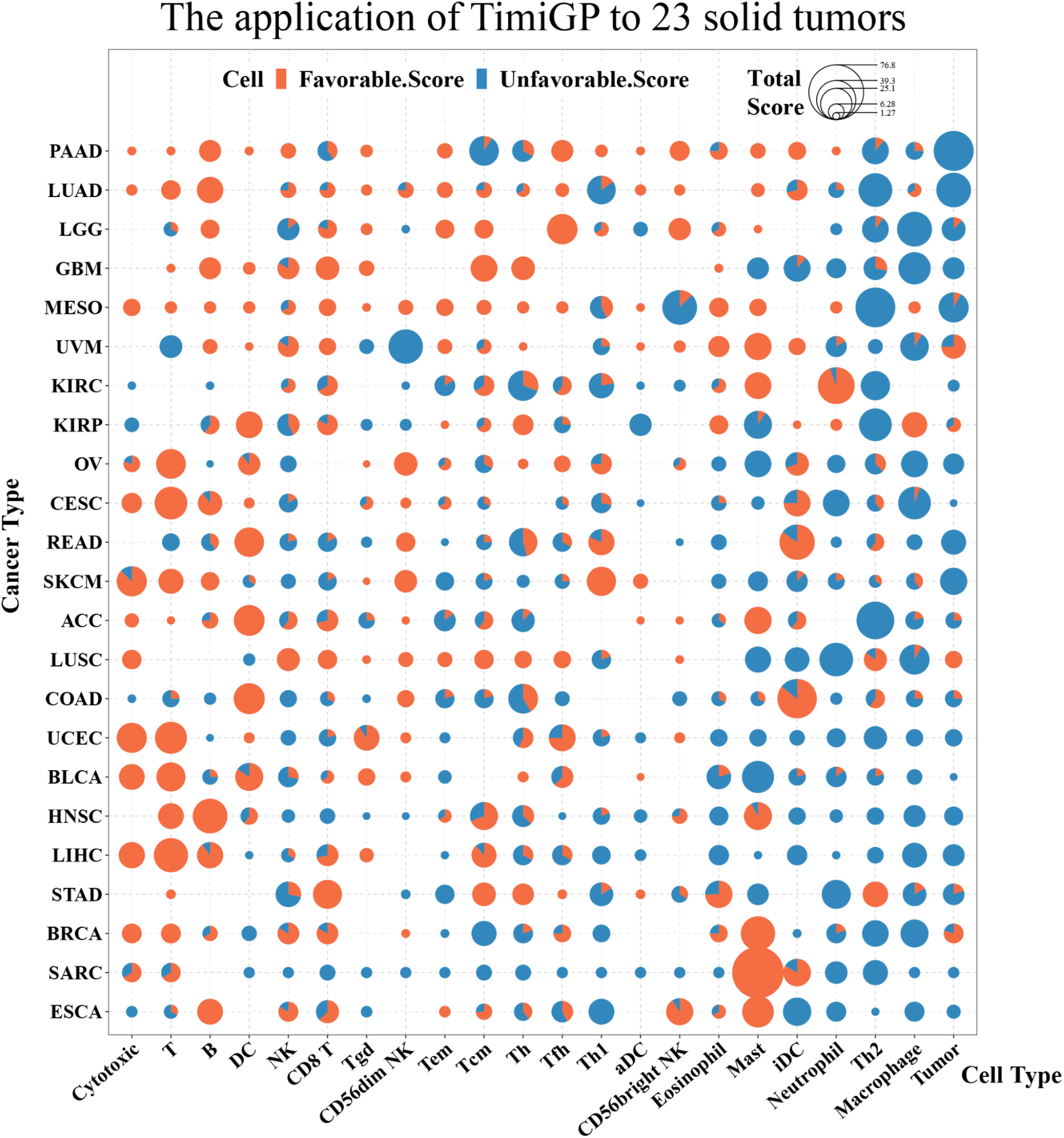
TimiGP reveals the prognostic value of immune cells in 23 solid tumors. Scatterpie chart showing TimiGP favorability score of cell types(x-axis) defined in modified immunoscore across 23 TCGA cancer types(y-axis). Each dot is a tiny pie chart showing the percentage of favorable(orange) or unfavorable(blue) scores. The area of the dot represents the sum of both scores. Abbreviation of cell types: B, B Cell; T, T cell; Th, Helper T cell; Th1, Type 1 Th; Th2, Type 2 Th; Tfh, Follicular Th; Tcm, Central memory T cell; Tem, Effector memory T cell; Tgd, Gamma delta T cell(Tγδ); Cytotoxic, Cytotoxic cell(common cytotoxic features of anti-tumor CD8 T cells, Tγδ, and NK cells); NK, Natural killer cell; DC, Dendritic cell; iDC, Immature DC; aDC, Activated DC; Mast, Mast cell; Tumor, Tumor cell. Abbreviation of cancer types: ACC, Adrenocortical carcinoma; BLCA, Bladder Urothelial Carcinoma; BRCA, Breast invasive carcinoma; CESC, Cervical squamous cell carcinoma and endocervical adenocarcinoma; COAD, Colon adenocarcinoma; ESCA, Esophageal carcinoma; GBM, Glioblastoma multiforme; HNSC, Head and Neck squamous cell carcinoma; KIRC, Kidney renal clear cell carcinoma; KIRP, Kidney renal papillary cell carcinoma; LGG, Brain Lower Grade Glioma; LIHC, Liver hepatocellular carcinoma; LUAD, Lung adenocarcinoma; LUSC, Lung squamous cell carcinoma; MESO, Mesothelioma; OV, Ovarian serous cystadenocarcinoma; PAAD, Pancreatic adenocarcinoma; READ, Rectum adenocarcinoma; SARC, Sarcoma; SKCM, Skin Cutaneous Melanoma; STAD, Stomach adenocarcinoma; UCEC, Uterine Corpus Endometrial Carcinoma; UVM, Uveal Melanoma. See also Figure S5.

Moreover, TimiGP also revealed the heterogeneity between distinct cancer types, in line with the previous studies^2, 7, 9^. Interestingly, pancreatic cancer, a typically immune “cold” tumor with low immune infiltration^67,68^, could pass our filter on immune marker expressions and be amenable for TimiGP analysis (Figure S5C, D). As expected, the immune cold pancreatic cancer exhibited distinct functional interaction networks compared to immune hot melanoma. Of note, although the interaction of cytotoxic cells → tumor cells, the preset controls, was identified as a functional interaction in pancreatic cancer, cytotoxic cells themselves were not identified as a predominant favorable cell type for anti-tumor immunity and prognosis. Instead, our analysis indicated that B cells, reported as salient features of pancreatic cancer^69–71^, might play a more important role in anti-cancer immunity in this cancer type, highlighting the variability in immune responses against different cancer types. Altogether, these results demonstrate that TimiGP is a promising method to analyze TIME interactions and identify prognostic cell types across different cancers.

## Discussion

In this study, we describe TimiGP as a new platform for decoding inter-cell functional interactions and the clinical values of immune cells from gene expression data. TimiGP is distinguished from related computational approaches in two aspects. First, unlike existing methods based on ligand-receptor interactions^28^, we studied the TIME from a different perspective - the potential relationships between immune cells’ functions. Referring to the dynamic balance of the immune system^1, 72^, we proposed the novel concept of functional interactions to reveal the relative capacity of immune cells in both biological behaviors and clinical values (Figure S6A). Second, compared to previous deconvolution methods^15–25^, TimiGP starts with pairwise relations at the expression level and estimates clinical values of immune cells through network analysis, which avoids the prognostic bias caused by immune cell co-infiltration. Therefore, TimiGP is not restricted by ligand-receptor knowledge or immune crosstalk influence and can identify cell target-to-hit and study the underlying causes of clinical heterogeneity.

TimiGP also paves the path to define comprehensive cell atlases (Figure S6B). Since it compares the paired gene expression of individual tumor samples, it does not require complicated normalization. This makes TimiGP applicable for integrative analysis with the transcriptomic profiles obtained from different specimens (e.g., fresh, frozen, FFPE) and sequencing platforms (e.g., RNA-seq, microarray). Furthermore, TimiGP is capable of the comprehensive assessment of the tumor microenvironment at different scales. It only depends on reliable cell-type markers and does not require complex training on gene signature expression references. This allows it to directly utilize cell-type markers obtained from scRNA-seq analysis and hence be flexible to achieve a higher resolution of cell subpopulations or a broader scale of the tumor microenvironment, such as the entire microenvironment, the TIME, and T cell subtypes as shown in our studies. Although the quality of cell-type markers remains a determining factor influencing the robustness of our method, the rapid pace of scRNA-seq makes this requirement unlikely to impede TimiGP application. In addition, even though we mainly demonstrate the utility of TimiGP in melanoma, it is generalizable to pan-cancer studies and other diseases for which expression profiles and clinical statistics are available.

Compared to predictors based on individual gene expressions, gene pair-based prognostic models have better normalizing robustness, predicting accuracy, and translational potential^73–76^. Despite this, it remains statistically and computationally challenging to identify the best combination of gene pair signatures due to the quadratically combinatorial complexity. While recent studies have revealed the advantage of immune-related gene pairs in developing prognostic signatures^36, 52^, the quadratic number of candidate genes still complicates the modeling process. Recent efforts focused on the improvement of machine-learning algorithms for variable selection and model fittings, which remain of poor quality in oncology^77^. Unlike those studies, TimiGP is an efficient algorithm to identify gene pairs representing functional inter-cell interactions, namely, a computational-friendly feature selector based on biological and clinical insights. Since TimiGP provides interpretable IMGP signatures by evaluating immune contexture, the corresponding prognostic model, TimiRS, can be validated across multiple independent datasets without complex model and training/testing/tuning processes. This suggests the necessity of understanding the TIME to develop prognostic models instead of just relying on computational modeling. In this study, we developed a simple prognostic model, TimiRS-Tem, based on effector memory CD8 T cells to showcase how TimiGP efficiently facilitates the development and endows high performance and interpretability to the prognostic model. Such TimiGP-based models are conducive to summarizing why and how predictors work in a certain cohort for further investigation, which will overcome another limitation of machine learning methods and expedite personalized therapy in cancer treatment. Altogether, TimiGP will offer substantial potential in clinical practice following further optimization.

In summary, TimiGP represents a broadly applicable framework to dissect the functional interactions and clinical values of infiltrating cells. This strategy can be used to investigate the clinically relevant tumor microenvironment at different scales and enables high-resolution analysis from low-resolution bulk profiles with the help of scRNA-seq markers. It systematically determines the associations between cellular biological functions and clinical outcomes through network analysis. Informed by such insights, TimiGP could therefore have utility for biological discovery and prediction. Given the method’s versatility (Figure S6B), we anticipate that TimiGP will be a useful asset for the practice of personalized medicine.

## Methods

### Datasets

The Cancer Genome Atlas (TCGA)^78^ RNA-seq dataset for 23 solid tumors (ACC, sBRCA, COAD, GBM, KIRC, LGG, LUAD, MESO, PAAD, SARC, STAD, UVM, BLCA, CESC, ESCA, HNSC, KIRP, LIHC, LUSC, OV, READ, SKCM, UCEC) was downloaded from Firehose (https://gdac.broadinstitute.org/). This dataset consisted of RSEM-normalized gene expression data for 20,501 genes. Of the SKCM samples, 368 are from metastatic tumor samples (TCGA_SKCM06)^39^. Among them, 357 samples with survival statistics for TimiGP analysis and prognostic model training. In addition, five independent microarray datasets (GSE65904^54^, GSE22155^55^, GSE54467^56^, GSE53118^57^, GSE8401^58^) for metastatic melanoma were used as validation.

### Immune infiltration estimation

The immune infiltration estimated by MCP-counter^17^, TIMER^18, 19^, CIBERSORT^22^, EPIC^24^, xCELL^15^, and quanTIseq^23^ was downloaded from TIMER2.0 website^20, 25^ (http://timer.cistrome.org/infiltration_estimation_for_tcga.csv.gz). The results of TCGA_SKCM06 cohorts were selected for this study. Of note, “Absolute Score” was used as the result of CIBERSORT to ensure the result is comparable across samples.

The results of CIBERSORTx^16^ (https://cibersortx.stanford.edu/) and EcoTyper^21^ (https://ecotyper.stanford.edu/carcinoma/) were calculated through their websites, whose input was the TPM normalized expression of TCGA_SKCM06 cohorts. According to the instructions with CIBERSORTx, we chose the LM22 signature matrix, disabled batch correction, absolute run mode, and 100 permutations. As for the cell states estimation, we ran EcoTyper with default parameters. Those inferred cell/state infiltrations were used for survival analysis (univariate Cox proportional hazards regression, Figure 1A, Table S1A) and co-infiltration analysis(Pearson correlation and hierarchical clustering, Figure S1).

### IMG’s prognostic value evaluation

The immune marker genes (IMGs) were collected from several studies^40–42^ and supplemented by well-known immune-related regulatory genes. Their prognostic values were analyzed by the univariate Cox regression (Figure 2B, Table S1B). The prognostic IMGs (BH-adjusted P-Value < 0.05) were further evaluated by Pearson correlation and hierarchical clustering to demonstrate the co-expression patterns (Figure 1C). The co-expression pattern of all collected IMGs and the cell-type markers used in MCP-counter, TIMER, CIBERSORT(x), EPIC, xCELL, quanTIseq are analyzed in the same way, respectively (Figure S1B).

### TimiGP Framework

TimiGP is designed to infer functional inter-cell interactions and estimate clinical values of immune cells based on transcriptomic profiles of bulk tissues. Therefore, the method’s inputs are immune marker gene expression profile and patient survival statistics (Figure 2).

#### Step 1: Pairwise Comparison

Under the hypothesis that the expression ratios between IMGs capture critical immune interactions, TimiGP defines marker pair score in the first step. After gene-wise median normalization, it captures the logical relation of any two selected genes at the expression level using the following expression:

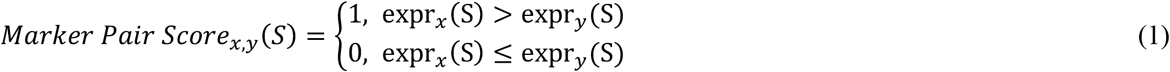

where *expr_x_*(*s*) and *expr_y_*(*s*) represent the expression of genes x and y in sample S.

By applying this function to gene pairs in each sample, TimiGP will generate a matrix of marker pair scores(MPS).

#### Step 2: Prognostic IMGP Selection

Based on MPS, TimiGP utilizes univariate Cox regression to select prognostic marker pairs. Since each gene pair has two directions, x →y and y →x, and the MPS is a logic value, the expression of x greater than y(MPS_x,y_=1) associated with favorable prognosis is the same as the expression of y greater than x(MPS_y,x_=1) associated with unfavorable prognosis. To remove the duplicates, we only consider the mark pair with an MPS of 1 associated with a good prognosis. Therefore, the marker pair with “MPS=1, Hazard Ratio < 1, BH-adjusted P-value < 0.05” are selected as prognostic IMGP.

#### Step 3: cell Interaction Annotation

Based on the characteristics of the immune system, we assume that the balance between pro- and anti-tumor immunity determines effective or suppressive TIME and subsequently impacts prognosis(Figure S6A). We simplify the complex TIME balance to the relative function between paired immune cells, whose functions are represented by their marker genes. Accordingly, TimiGP will generate an annotation set that includes all potential cell-cell interactions and is denoted by marker pairs. In the annotation, a cell interaction X →Y is defined by a list of marker pairs that points from the markers of cell X (e.g., x1, x2) to the markers of cell Y(e.g., y1, y2) as below:

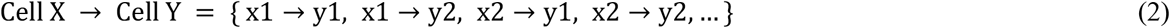

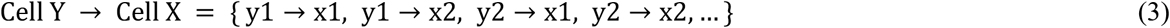

#### Step 4: Enrichment Analysis

To determine which cell interactions are associated with a favorable prognosis, TimiGP will perform the enrichment analysis. It statistically examines whether a cell interaction is over-represented in the prognostic pairs by calculating the enrichment ratio and p-value based on hypergeometric distribution^79^:

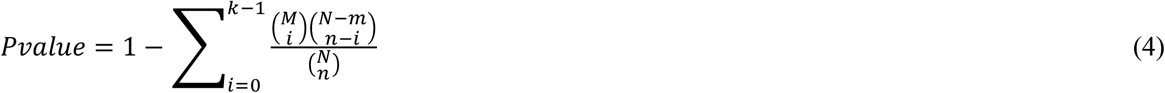

Where N is the total number of annotated gene pairs including both directions, such as x→y and y→x, M is the number of gene pairs of specific cell interaction, n is the size of prognostic gene pairs, and k is the number of prognostic gene pairs annotated by the cell interaction.

If the marker pairs of a cell interaction are significantly enriched in the prognostic pairs (BH-adjusted P-value < 0.05), the cell interaction is characterized as a “functional interaction”, which represents relative capacity between two cells about both biological and clinical roles. For example, functional cell X→Y interaction denotes that the more effective X function than Y positively impacts survival. Cell X is an anti-tumor cell associated with a favorable prognosis, while cell Y is likely to be a pro-tumor cell type associated with an inferior outcome.

#### Step 5: Network Analysis

By integrating all statistically significant enrichment results, TimiGP will construct a functional interaction network at the cell level, which provides a complete portrait of the TIME. In the network, the node represents the cell type, and the edge represents the functional interaction described. Given that the ingoing and outgoing edges demonstrate opposite functions of the cells, the relative function of involved cell types is scored according to the degrees, called the favorability score. Considering the dual roles of immune cells, we calculated the favorability score composed of favorable and unfavorable scores as follows:

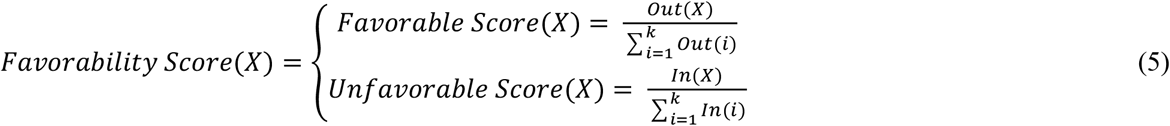

In the equation, Out(X) and In(X) represent the number of out-degree and in-degree for node cell X, respectively. The favorability score evaluates the relative role of cell X among k cell types in the analysis. If the cell type has a higher favorable score, it is a potential anti-tumor cell associated with an improved prognosis. If it has a higher unfavorable score, it is more likely to play a pro-tumor role associated with poor prognosis than other cells. If it has both higher favorable and unfavorable scores, we should come back to examine the network to figure out which cell type it is more or less favorable than.

### Feature selection for prognostic model

Feature selection is a critical step in biomarker expression-based models. Since we aimed to demonstrate the application of TimiGP in this area rather than develop a clinical prognostic model, we referred to the current gene pair-based model^36^ and utilized regularized cox model with the L1 penalty (least absolute shrinkage and selection operator, LASSO) to select features(glmnet R package v4.1-2)^53, 80^. The penalty parameter was estimated by 10-fold cross-validation and measured by Harrel C-index. We chose “lambda.1se” as the value of λ. To evaluate the robustness of the selected IRGPs, feature selection was repeated 1000 times in 80% randomized TCGA_SKCM06 cohorts. Since the cell-type markers summarized by Charoentong, P. et al.^41^ have the largest number of mutually exclusive genes in this study, we chose their expression profiles as the input for further analysis(Figure 4). To prove TimiGP can benefit feature selection, we compared the probability of the same IMGP selected from prognostic and enriched pairs by LASSO. Notably, prognostic pairs were obtained from the second step of TimiGP as control, which was used by current methods, and enriched pairs were from the fourth step, representing the functional interactions(Figure 2). The probability that marker pair x→y is selected as a predictor will be calculated below(Figure 4B):

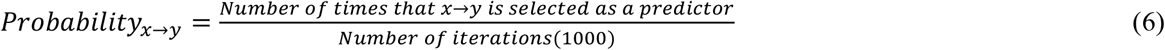

To examine if TimiGP can facilitate the prognostic model development, we next selected the top 21 IMGPs following the selection probability for prognostic models. The number of IMGPs was determined by 10-fold cross-validation according to C-index in the TCGA_SKCM06 training set. The signature for the control model was picked from prognostic pairs, and the signature for the Tem model was selected from enriched pairs that represent Effector memory CD8 T cell (CD8 Tem)-to-other cells interactions with strong confidence(Adjust.P.Value < 0.0001).

### TimiRS model construction and validation

Since the gene pair represents the intuitive notion that its high relative abundance is associated with a good prognosis, the risk score (TimiRS), on the contrary, is calculated by the percentage of predictors whose marker pair scores are 0.

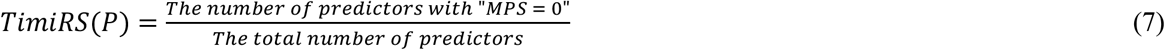

We considered TCGA_SKCM06 as a training set because the signature was selected based on the cohorts and validated the models in five independent datasets (GSE65904^54^, GSE22155^55^, GSE54467^56^, GSE53118^57^, GSE8401^58^). Among 141 functional interactions identified by TimiGP, each interaction has a list of supportive enriched pairs. They were applied as individual models to calculate TimiRS in training and five validation sets. The prognostic model of these signatures was evaluated by Cox regression which fits TimiRS as a continuous variable(P-Value < 0.05, Figure 4C, D).

As for TimiRS-control and TimiRS-Tem models, we used the signatures selected from prognostic pairs (control) or enriched pairs related to CD8 Tem →other cells interactions (Tem) to calculate TimiRS in training and five independent datasets. The performance of each model was evaluated by the C-index, the ability of patient stratification (Kaplan-Meier estimate using the median as a cutoff), and time-dependent receiver operating characteristic (ROC).

### Pan-cancer TimiGP analysis

Considering the quality of survival statistics, we selected 23 solid tumors (ACC, BRCA, COAD, GBM, KIRC, LGG, LUAD, MESO, PAAD, SARC, STAD, UVM, BLCA, CESC, ESCA, HNSC, KIRP, LIHC, LUSC, OV, READ, SKCM, UCEC) from TCGA for pan-cancer analysis^78^. All samples with the survival information were used to perform TimiGP analysis per cancer type. To be consistent across 23 tumor types, we selected the top 4352 prognostic IMGPs following P-value in step 2 for each cancer type. The number was determined by the cancer type with the least number of prognostic IMGPs (P-value < 0.05). We visualized the TimiGP results by scatterpie R package (v0.1.6) (Figure 6). We also selected the top 10 functional interactions related to Cytotoxic cell →Other cells or Other cells →Tumor cells interactions ranked by the number of cancer types from which the interactions were identified.

### Statistical analysis

The hierarchical network in Figure 3C was visualized by Cytoscape (v3.9.0)^81^ and structured by yFile with default parameters. Statistical analyses were performed with R (v4.1.0) and visualized with ggplot2 R package (v3.3.5)^82^, scatterpie R package (v0.1.6)^83^, VennDiagram R package (v1.6.20))^84^, ComplexHeatmap R package (v2.10.0)^85^, and circlize R package (v0.4.13)^86^. If not mentioned in the specific method section, the cox regression and C-index were performed using survival R package (v3.2-11)^87^. Time-dependent ROC at 3 years and the corresponding AUC(Area under the ROC Curve) were calculated by survivalROC R package (v1.0.3)^88^. The Kaplan-Meier estimate was performed through survminer R package (v0.4.9)^89^.

## Data availability

All bulk RNA-seq and microarray data and cell type markers are publicly available. The modified cell-type annotations of Bindea2013(immunoscore)^40^, Charoentong2017^41^, Xu2018(TIP)^42^, Newman2015(LM22)^22^, Tirosh2016(metastatic melanoma scRNA-seq)^64^, Zheng2021(pan-cancer T cell scRNA-seq)^65^ are incorporated in TimiGP R package (https://github.com/CSkylarL/TimiGP).

## Code availability

The original code for TimiGP is publicly available as of the publication date for non-profit academic use. The TimiGP R package and updates are available via GitHub: https://github.com/CSkylarL/TimiGP. All code in this study is deposited in GitHub: https://github.com/CSkylarL/MSofTimiGP.

## Acknowledgments

This work was supported by the National Cancer Institute of the National Institute of Health Research Project Grant (J.Z., R01CA234629-01), the AACR-Johnson & Johnson Lung Cancer Innovation Science Grant (J.Z., 18-90-52-ZHAN), the MD Anderson Physician Scientist Program(J.Z.), the MD Anderson Lung Cancer Moon Shot Program(J.Z.) and the Cancer Prevention Research Institute of Texas (CPRIT) (C.C., RR180061). C.C. is a CPRIT Scholar in Cancer Research. The authors would like to acknowledge the support of the High Performance Computing for research facility at the University of Texas MD Anderson Cancer Center for providing computational resources that have contributed to the research results reported in this paper. The Figure 1E, Figure 2, Figure 4A, Figure S6 were created with BioRender.com.

## Author contributions

Conceptualization, C.L. and C.C.; Methodology, C.L. and C.C.; Software: C.L.; Validation: C.L.; Formal Analysis: C.L.; Investigation, C.L., B.Z. and E.S.; Resources: C.C and J.Z.; Data Curation: C.C and C.L; Writing – Original Draft, C.L.; Writing – Review & Editing, J.Z., C.C, A.R., B.Z. and E.S.; Visualization, C.L.; Supervision: J.Z. and C.C.; Funding Acquisition, J.Z. and C.C..

## Corresponding authors

Correspondence to Jianjun Zhang and Chao Cheng

## Declaration of interests

A.R. serves on the Scientific Advisory Board and has received honoraria from Adaptive Biotechnologies. J.Z. reports grants from Merck, grants and personal fees from Johnson and Johnson and Novartis, personal fees from Bristol Myers Squibb, AstraZeneca, GenePlus, Innovent and Hengrui outside the submitted work. The remaining authors declare no potential conflicts of interest.

## Supplemental information

**Figure S1** The inter-cell co-infiltration and cell-type marker co-expression in metastatic melanoma, related to Figure 1

**Figure S2** Examples that pairwise relations between gene expressions reduce the prognostic bias, related to Figure 1

**Figure S3** Evaluation of TimiGP robustness with distinct cell type annotations, related to Figure 3

**Figure S4** Better performance of the TimiRS-Tem model than TimiRS-Control model in independent validation sets, related to Figure 4

**Figure S5** Pan-cancer functional interactions related to control cells and TimiGP analysis of pancreatic adenocarcinoma, related to Figure 6

**Figure S6** Summary of TimiGP Rationale and Applications, related to Discussion

**Table S1** Prognostic Value of Immune cell infiltration and Immune marker gene expressions

(A) Prognostic Value of Immune cell infiltration estimated by existing methods.

(B) Prognostic Value of Immune marker genes collected for this study.

**Table S3** TimiGP output with Bindea2013 cell type annotation for metastatic melanoma

**Table S4** Probability of IMGPs selected from enriched or prognostic candidates by LASSO

**Table S5** Prognostic value of functional interactions

**Table S6** Model information about TimiRS-Tem and TimiRS-Control model

**Table S7** Overview of Survival Statistics of TCGA datasets

## Notes

https://github.com/CSkylarL/TimiGP

https://github.com/CSkylarL/MSofTimiGP

### Competing Interest Statement

Alexandre Reuben serves on the Scientific Advisory Board and has received honoraria from Adaptive Biotechnologies. Jianjun Zhang reports grants from Merck, grants and personal fees from Johnson and Johnson and Novartis, personal fees from Bristol Myers Squibb, AstraZeneca, GenePlus, Innovent and Hengrui outside the submitted work. The remaining authors declare no potential conflicts of interest.

## Reference

1. Dunn, G.P., Bruce, A.T., Ikeda, H., Old, L.J. & Schreiber, R.D. Cancer immunoediting: from immunosurveillance to tumor escape. Nature immunology 3, 991–998 (2002).

2. Galon, J. & Bruni, D. Tumor immunology and tumor evolution: intertwined histories. Immunity 52, 55–81 (2020).

3. Raskov, H., Orhan, A., Christensen, J.P. & Gögenur, I. Cytotoxic CD8+ T cells in cancer and cancer immunotherapy. British journal of cancer 124, 359–367 (2021).

4. Shimasaki, N., Jain, A. & Campana, D. NK cells for cancer immunotherapy. Nature reviews Drug discovery 19, 200–218 (2020).

5. Togashi, Y., Shitara, K. & Nishikawa, H. Regulatory T cells in cancer immunosuppression— implications for anticancer therapy. Nature reviews Clinical oncology 16, 356–371 (2019).

6. Veglia, F., Sanseviero, E. & Gabrilovich, D.I. Myeloid-derived suppressor cells in the era of increasing myeloid cell diversity. Nature Reviews Immunology 21, 485–498 (2021).

7. Fridman, W.H., Zitvogel, L., Sautès–Fridman, C. & Kroemer, G. The immune contexture in cancer prognosis and treatment. Nature reviews Clinical oncology 14, 717–734 (2017).

8. Binnewies, M. et al. Understanding the tumor immune microenvironment (TIME) for effective therapy. Nature medicine 24, 541–550 (2018).

9. Bruni, D., Angell, H.K. & Galon, J. The immune contexture and Immunoscore in cancer prognosis and therapeutic efficacy. Nature Reviews Cancer 20, 662–680 (2020).

10. Galon, J. & Bruni, D. Approaches to treat immune hot, altered and cold tumours with combination immunotherapies. Nature reviews Drug discovery 18, 197–218 (2019).

11. Wagner, A., Regev, A. & Yosef, N. Revealing the vectors of cellular identity with single-cell genomics. Nature biotechnology 34, 1145–1160 (2016).

12. Lähnemann, D. et al. Eleven grand challenges in single-cell data science. Genome biology 21, 1–35 (2020).

13. Marco-Puche, G., Lois, S., Benítez, J. & Trivino, J.C. RNA-Seq perspectives to improve clinical diagnosis. Frontiers in genetics 10, 1152 (2019).

14. Kuksin, M. et al. Applications of single-cell and bulk RNA sequencing in onco-immunology. European Journal of Cancer 149, 193–210 (2021).

15. Aran, D., Hu, Z. & Butte, A.J. xCell: digitally portraying the tissue cellular heterogeneity landscape. Genome biology 18, 1–14 (2017).

16. Newman, A.M. et al. Determining cell type abundance and expression from bulk tissues with digital cytometry. Nature biotechnology 37, 773–782 (2019).

17. Becht, E. et al. Estimating the population abundance of tissue-infiltrating immune and stromal cell populations using gene expression. Genome biology 17, 1–20 (2016).

18. Li, B. et al. Comprehensive analyses of tumor immunity: implications for cancer immunotherapy. Genome biology 17, 1–16 (2016).

19. Li, T. et al. TIMER: a web server for comprehensive analysis of tumor-infiltrating immune cells. Cancer research 77, e108–e110 (2017).

20. Li, T. et al. TIMER2. 0 for analysis of tumor-infiltrating immune cells. Nucleic acids research 48, W509–W514 (2020).

21. Luca, B.A. et al. Atlas of clinically distinct cell states and ecosystems across human solid tumors. Cell 184, 5482–5496. e5428 (2021).

22. Newman, A.M. et al. Robust enumeration of cell subsets from tissue expression profiles. Nature methods 12, 453–457 (2015).

23. Finotello, F. et al. Molecular and pharmacological modulators of the tumor immune contexture revealed by deconvolution of RNA-seq data. Genome medicine 11, 1–20 (2019).

24. Racle, J., de Jonge, K., Baumgaertner, P., Speiser, D.E. & Gfeller, D. Simultaneous enumeration of cancer and immune cell types from bulk tumor gene expression data. elife 6 (2017).

25. Sturm, G. et al. Comprehensive evaluation of transcriptome-based cell-type quantification methods for immuno-oncology. Bioinformatics 35, i436–i445 (2019).

26. Choi, H. et al. Transcriptome analysis of individual stromal cell populations identifies stroma-tumor crosstalk in mouse lung cancer model. Cell reports 10, 1187–1201 (2015).

27. Noёl, F. et al. Dissection of intercellular communication using the transcriptome-based framework ICELLNET. Nature communications 12, 1–16 (2021).

28. Armingol, E., Officer, A., Harismendy, O. & Lewis, N.E. Deciphering cell–cell interactions and communication from gene expression. Nature Reviews Genetics 22, 71–88 (2021).

29. Varn, F.S., Wang, Y., Mullins, D.W., Fiering, S. & Cheng, C. Systematic Pan-Cancer Analysis Reveals Immune Cell Interactions in the Tumor MicroenvironmentPan-Cancer Analysis of Immune Cell Interactions. Cancer research 77, 1271–1282 (2017).

30. Kluger, H.M. et al. Characterization of PD-L1 expression and associated T-cell infiltrates in metastatic melanoma samples from variable anatomic sites. Clinical Cancer Research 21, 3052–3060 (2015).

31. Li, G., Zhu, X. & Liu, C. Characterization of Immune Infiltration and Construction of a Prediction Model for Overall Survival in Melanoma Patients. Frontiers in oncology 11, 976 (2021).

32. Masucci, M.T., Minopoli, M. & Carriero, M.V. Tumor associated neutrophils. Their role in tumorigenesis, metastasis, prognosis and therapy. Frontiers in oncology 9, 1146 (2019).

33. Zhou, B., Lawrence, T. & Liang, Y. The role of plasmacytoid dendritic cells in cancers. Frontiers in Immunology, 4414 (2021).

34. Zhai, L. et al. IDO1 in cancer: a Gemini of immune checkpoints. Cellular & molecular immunology 15, 447–457 (2018).

35. Qin, S. et al. Novel immune checkpoint targets: moving beyond PD-1 and CTLA-4. Molecular cancer 18, 1–14 (2019).

36. Li, B., Cui, Y., Diehn, M. & Li, R. Development and validation of an individualized immune prognostic signature in early-stage nonsquamous non–small cell lung cancer. JAMA oncology 3, 1529–1537 (2017).

37. Auslander, N. et al. Robust prediction of response to immune checkpoint blockade therapy in metastatic melanoma. Nature medicine 24, 1545–1549 (2018).

38. Jorgovanovic, D., Song, M., Wang, L. & Zhang, Y. Roles of IFN-γ in tumor progression and regression: A review. Biomarker research 8, 1–16 (2020).

39. Akbani, R. et al. Genomic classification of cutaneous melanoma. Cell 161, 1681–1696 (2015).

40. Bindea, G. et al. Spatiotemporal dynamics of intratumoral immune cells reveal the immune landscape in human cancer. Immunity 39, 782–795 (2013).

41. Charoentong, P. et al. Pan-cancer immunogenomic analyses reveal genotype-immunophenotype relationships and predictors of response to checkpoint blockade. Cell reports 18, 248–262 (2017).

42. Xu, L. et al. TIP: A Web Server for Resolving Tumor Immunophenotype ProfilingTIP: Tracking Tumor Immunophenotype. Cancer research 78, 6575–6580 (2018).

43. Dudek, A.M., Martin, S., Garg, A.D. & Agostinis, P. Immature, semi-mature, and fully mature dendritic cells: toward a DC-cancer cells interface that augments anticancer immunity. Frontiers in immunology 4, 438 (2013).

44. Murphy, K. & Weaver, C. Janeway’s immunobiology. (Garland science, 2016).

45. Komi, D.E.A. & Redegeld, F.A. Role of mast cells in shaping the tumor microenvironment. Clinical reviews in allergy & immunology 58, 313–325 (2020).

46. Cózar, B. et al. Tumor-Infiltrating Natural Killer CellsTumor-infiltrating Natural Killer Cells. Cancer discovery 11, 34–44 (2021).

47. O’Dwyer, R.T. et al. (American Society of Clinical Oncology, 2019).

48. Cohen, J.T., Miner, T.J. & Vezeridis, M.P. Is the neutrophil-to-lymphocyte ratio a useful prognostic indicator in melanoma patients? Melanoma Management 7, MMT47 (2020).

49. Bartlett, E.K. et al. High neutrophil-to-lymphocyte ratio (NLR) is associated with treatment failure and death in patients who have melanoma treated with PD-1 inhibitor monotherapy. Cancer 126, 76–85 (2020).

50. Capone, M. et al. Baseline neutrophil-to-lymphocyte ratio (NLR) and derived NLR could predict overall survival in patients with advanced melanoma treated with nivolumab. Journal for immunotherapy of cancer 6, 1–7 (2018).

51. Ma, J. et al. Neutrophil-to-lymphocyte Ratio (NLR) as a predictor for recurrence in patients with stage III melanoma. Scientific reports 8, 1–6 (2018).

52. Huang, R.-z. et al. Development of an immune-related gene pairs index for the prognosis analysis of metastatic melanoma. Scientific Reports 11, 1–12 (2021).

53. Simon, N., Friedman, J., Hastie, T. & Tibshirani, R. Regularization paths for Cox’s proportional hazards model via coordinate descent. Journal of statistical software 39, 1 (2011).

54. Cirenajwis, H. et al. Molecular stratification of metastatic melanoma using gene expression profiling: Prediction of survival outcome and benefit from molecular targeted therapy. Oncotarget 6, 12297 (2015).

55. Jönsson, G. et al. Gene expression profiling–based identification of molecular subtypes in stage IV melanomas with different clinical outcome. Clinical cancer research 16, 3356–3367 (2010).

56. Jayawardana, K. et al. Determination of prognosis in metastatic melanoma through integration of clinico-pathologic, mutation, mRNA, microRNA, and protein information. International journal of cancer 136, 863–874 (2015).

57. Mann, G.J. et al. BRAF mutation, NRAS mutation, and the absence of an immune-related expressed gene profile predict poor outcome in patients with stage III melanoma. Journal of Investigative Dermatology 133, 509–517 (2013).

58. Xu, L. et al. Gene expression changes in an animal melanoma model correlate with aggressiveness of human melanoma metastases. Molecular Cancer Research 6, 760–769 (2008).

59. Zhang, P., Côté, A.L., de Vries, V.C., Usherwood, E.J. & Turk, M.J. Induction of postsurgical tumor immunity and T-cell memory by a poorly immunogenic tumor. Cancer research 67, 6468–6476 (2007).

60. Wistuba-Hamprecht, K. et al. Peripheral CD8 effector-memory type 1 T-cells correlate with outcome in ipilimumab-treated stage IV melanoma patients. European Journal of Cancer 73, 61–70 (2017).

61. Svensson, V., Vento-Tormo, R. & Teichmann, S.A. Exponential scaling of single-cell RNA-seq in the past decade. Nature protocols 13, 599–604 (2018).

62. Papalexi, E. & Satija, R. Single-cell RNA sequencing to explore immune cell heterogeneity. Nature Reviews Immunology 18, 35–45 (2018).

63. Turley, S.J., Cremasco, V. & Astarita, J.L. Immunological hallmarks of stromal cells in the tumour microenvironment. Nature reviews immunology 15, 669–682 (2015).

64. Tirosh, I. et al. Dissecting the multicellular ecosystem of metastatic melanoma by single-cell RNA-seq. Science 352, 189–196 (2016).

65. Zheng, L. et al. Pan-cancer single-cell landscape of tumor-infiltrating T cells. Science 374, abe6474 (2021).

66. Miller, B.C. et al. Subsets of exhausted CD8+ T cells differentially mediate tumor control and respond to checkpoint blockade. Nature immunology 20, 326–336 (2019).

67. Kleeff, J. et al. Pancreatic cancer. Nature reviews Disease primers 2, 1–22 (2016).

68. Hilmi, M., Bartholin, L. & Neuzillet, C. Immune therapies in pancreatic ductal adenocarcinoma: Where are we now? World journal of gastroenterology 24, 2137 (2018).

69. Spear, S. et al. Discrepancies in the tumor microenvironment of spontaneous and orthotopic murine models of pancreatic cancer uncover a new immunostimulatory phenotype for B cells. Frontiers in immunology 10, 542 (2019).

70. Castino, G.F. et al. Spatial distribution of B cells predicts prognosis in human pancreatic adenocarcinoma. Oncoimmunology 5, e1085147 (2016).

71. Tewari, N. et al. The presence of tumour-associated lymphocytes confers a good prognosis in pancreatic ductal adenocarcinoma: an immunohistochemical study of tissue microarrays. BMC cancer 13, 1–10 (2013).

72. Cicchese, J.M. et al. Dynamic balance of pro-and anti-inflammatory signals controls disease and limits pathology. Immunological reviews 285, 147–167 (2018).

73. Donald, G. & Christian, d.A. Classifying gene expression profiles from pairwise mRNA comparisons. Statistical applications in genetics and molecular biology 3, 1–22 (2004).

74. Leek, J.T. The tspair package for finding top scoring pair classifiers in R. Bioinformatics 25, 1203–1204 (2009).

75. Patil, P., Bachant-Winner, P.-O., Haibe-Kains, B. & Leek, J.T. Test set bias affects reproducibility of gene signatures. Bioinformatics 31, 2318–2323 (2015).

76. Shen, R., Luo, L. & Jiang, H. Identification of gene pairs through penalized regression subject to constraints. BMC bioinformatics 18, 1–11 (2017).

77. Dhiman, P. et al. Risk of bias of prognostic models developed using machine learning: a systematic review in oncology. Diagn Progn Res 6, 13 (2022).

78. Weinstein, J.N. et al. The cancer genome atlas pan-cancer analysis project. Nature genetics 45, 1113–1120 (2013).

79. Boyle, E.I. et al. GO:: TermFinder—open source software for accessing Gene Ontology information and finding significantly enriched Gene Ontology terms associated with a list of genes. Bioinformatics 20, 3710–3715 (2004).

80. Friedman, J., Hastie, T. & Tibshirani, R. Regularization paths for generalized linear models via coordinate descent. Journal of statistical software 33, 1 (2010).

81. Shannon, P. et al. Cytoscape: a software environment for integrated models of biomolecular interaction networks. Genome research 13, 2498–2504 (2003).

82. Wickham, H. Package ‘ggplot2’: elegant graphics for data analysis. Springer-Verlag New York. doi 10, 978–970 (2016).

83. Guangchuang, Y. (2021).

84. Chen, H. VennDiagram: generate high-resolution Venn and Euler plots. R package version 1.6. 20. Website https://CRAN.R-project.org/package= VennDiagram (2018).

85. Gu, Z., Eils, R. & Schlesner, M. Complex heatmaps reveal patterns and correlations in multidimensional genomic data. Bioinformatics 32, 2847–2849 (2016).

86. Gu, Z., Gu, L., Eils, R., Schlesner, M. & Brors, B. Circlize implements and enhances circular visualization in R. Bioinformatics 30, 2811–2812 (2014).

87. Therneau, T.M. & Grambsch, P.M. in Modeling survival data: extending the Cox model 39–77 (Springer, 2000).

88. Heagerty, P.J., Saha-Chaudhuri, P. & Saha-Chaudhuri, M.P. Package ‘survivalROC’. San Francisco: GitHub (2013).

89. Kassambara, A., Kosinski, M., Biecek, P. & Fabian, S. Survminer: Drawing Survival Curves Using Ggplot2. 2021. URL https://CRAN.R-project.org/package=survminer. R package version 0.4 9 (2021).

