## Supplementary_Figure for "TimiGP: inferring inter-cell functional interactions and clinical values in the tumor immune microenvironment through gene pairs"

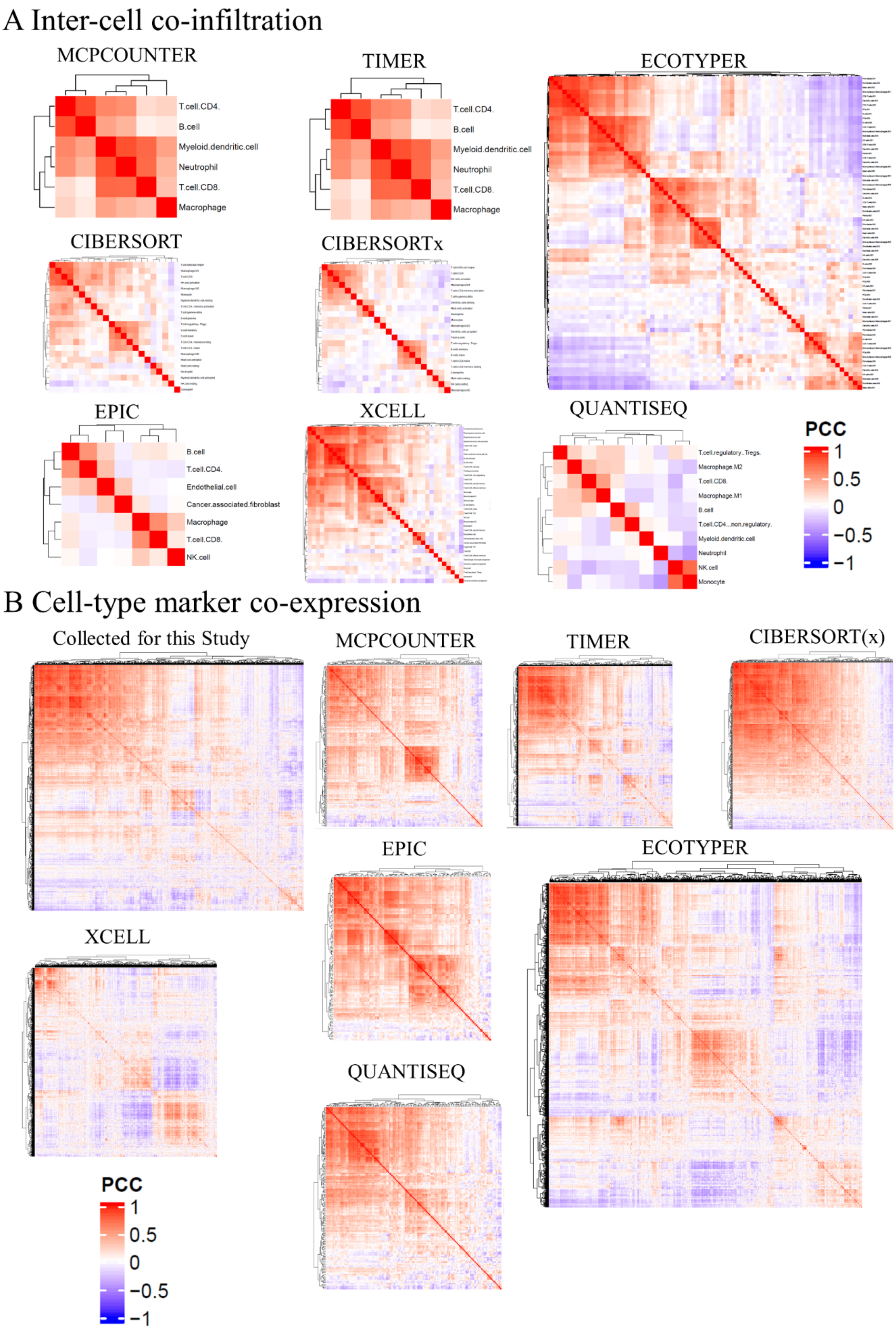
A Inter-cell co-infiltration
MCPCOUNTER
T.cell.CD4.
B.cell
Myeloid.dendritic.cell
Neutrophil
T.cell.CD8.
Macrophage
TIMER
T.cell.CD4.
B.cell
Myeloid.dendritic.cell
Neutrophil
T.cell.CD8.
Macrophage
ECOTYPER
CIBERSORT
CIBERSORTx
EPIC
B.cell
T.cell.CD4.
Endothelial.cell
Cancer.associated.fibroblast
Macrophage
T.cell.CD8.
NK.cell
XCELL
QUANTISEQ
T.cell.regulatory..Tregs.
Macrophage.M2
T.cell.CD8.
Macrophage.M1
B.cell
T.cell.CD4...non.regulatory.
Myeloid.dendritic.cell
Neutrophil
NK.cell
Monocyte
PCC
1
0.5
0
−0.5
−1
B Cell-type marker co-expression
Collected for this Study
MCPCOUNTER
TIMER
CIBERSORT(x)
EPIC
ECOTYPER
XCELL
QUANTISEQ
PCC
1
0.5
0
−0.5
−1

### Figure S1 The inter-cell co-infiltration and cell-type marker co-expression in metastatic melanoma, related to Figure 1

(A) Heatmap of Pearson correlation coefficients(PCC) showing the inter-cell co-infiltration. The cell types inferred by MCP-counter, TIMER, EcoTyper, CIBERSORT, CIBERSORTx, EPIC, xCELL, or quanTIseq.

(B) Heatmap of Pearson correlation coefficients(PCC) showing the cell-type marker co-expression, There cell-type markers were collected for this study or used by the transcriptome-based cell-type quantification methods mentioned above. All analysis has been performed with the RNA-seq of metastatic melanoma in the Cancer Genome Atlas(TCGA_SKCM06).

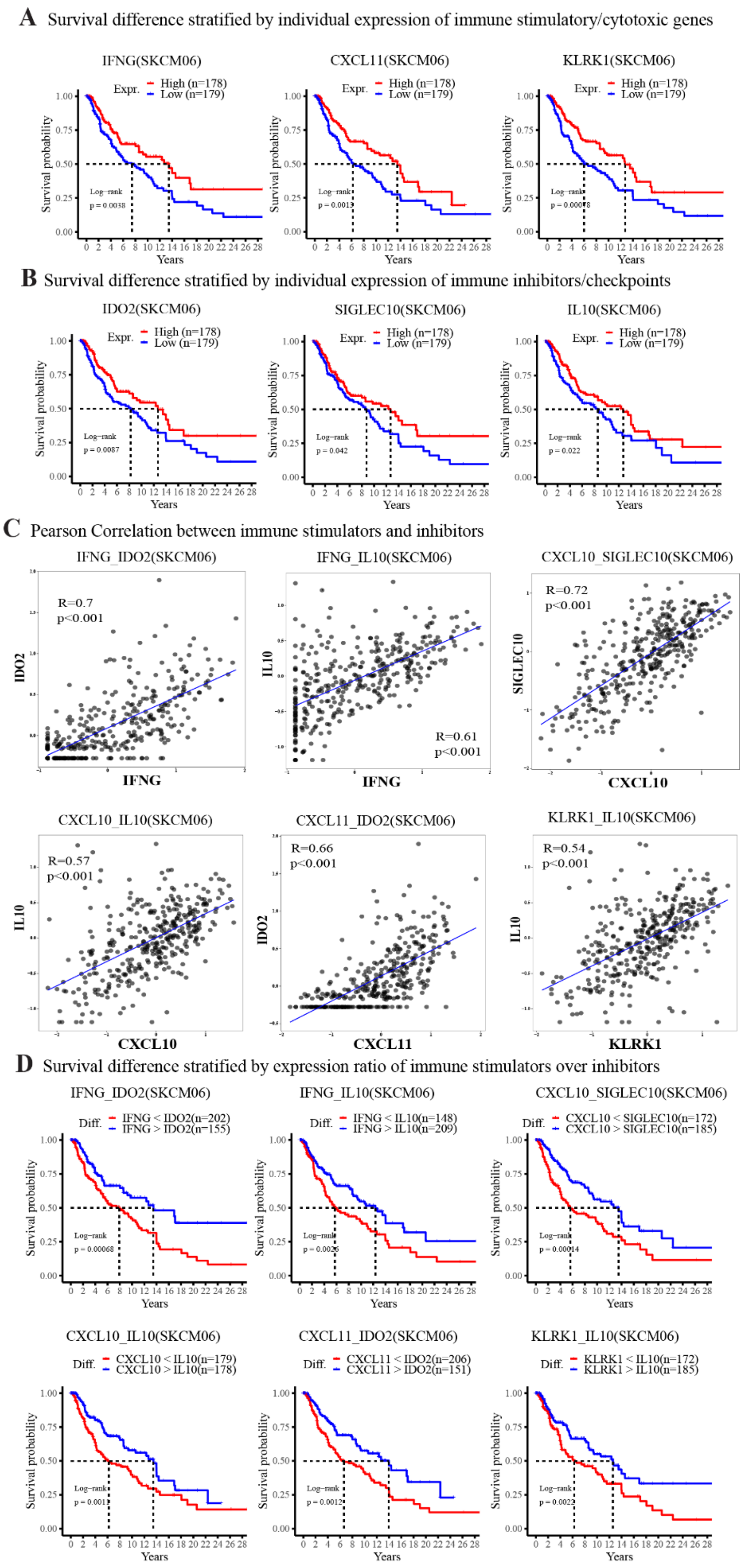
A Survival difference stratified by individual expression of immune stimulatory/cytotoxic genes
IFNG(SKCM06)
CXCL11(SKCM06)
KLRK1(SKCM06)
Expr.
High (n=178)
Low (n=179)
Survival probability
Years
B Survival difference stratified by individual expression of immune inhibitors/checkpoints
IDO2(SKCM06)
SIGLEC10(SKCM06)
IL10(SKCM06)
C Pearson Correlation between immune stimulators and inhibitors
IFNG_IDO2(SKCM06)
R=0.7
p<0.001
IFNG_IL10(SKCM06)
R=0.61
p<0.001
CXCL10_SIGLEC10(SKCM06)
R=0.72
p<0.001
CXCL10_IL10(SKCM06)
R=0.57
p<0.001
CXCL11_IDO2(SKCM06)
R=0.66
p<0.001
KLRK1_IL10(SKCM06)
R=0.54
p<0.001
D Survival difference stratified by expression ratio of immune stimulators over inhibitors
IFNG_IDO2(SKCM06)
Diff.
IFNG < IDO2(n=202)
IFNG > IDO2(n=155)
IFNG_IL10(SKCM06)
IFNG < IL10(n=148)
IFNG > IL10(n=209)
CXCL10_SIGLEC10(SKCM06)
CXCL10 < SIGLEC10(n=172)
CXCL10 > SIGLEC10(n=185)
CXCL10_IL10(SKCM06)
CXCL10 < IL10(n=179)
CXCL10 > IL10(n=178)
CXCL11_IDO2(SKCM06)
CXCL11 < IDO2(n=206)
CXCL11 > IDO2(n=151)
KLRK1_IL10(SKCM06)
KLRK1 < IL10(n=172)
KLRK1 > IL10(n=185)

### Figure S2 Examples that pairwise relations between gene expressions reduce the prognostic bias, related to Figure 1

(A) Kaplan-Meier(KM) curve showing the overall survival of patients with high(red) or low(blue) expression of immune stimulatory or cytotoxic genes: IFNG(left), CXCL11(middle), or KLRK1(right) using the median as a cutoff.

(B) KM curve showing the overall survival of patients with high(red) or low(blue) expression of immune inhibitors or checkpoints: IDO2(left), SIGLEC10(middle), or IL10(right) using the median as a cutoff.

(C) Scatter plot showing the Pearson correlation between immune effector and suppressor genes.

(D) KM curve showing the overall survival of patients with different ratios of immune effector-over-suppressor genes. The red represents the expression of immune effectors less than suppressor genes, and the blue shows the opposite group.

All analysis has been performed with the RNA-seq of metastatic melanoma in the Cancer Genome Atlas(TCGA_SKCM06). Significance on KM plot is calculated by the log-rank test comparing the survival of two groups.

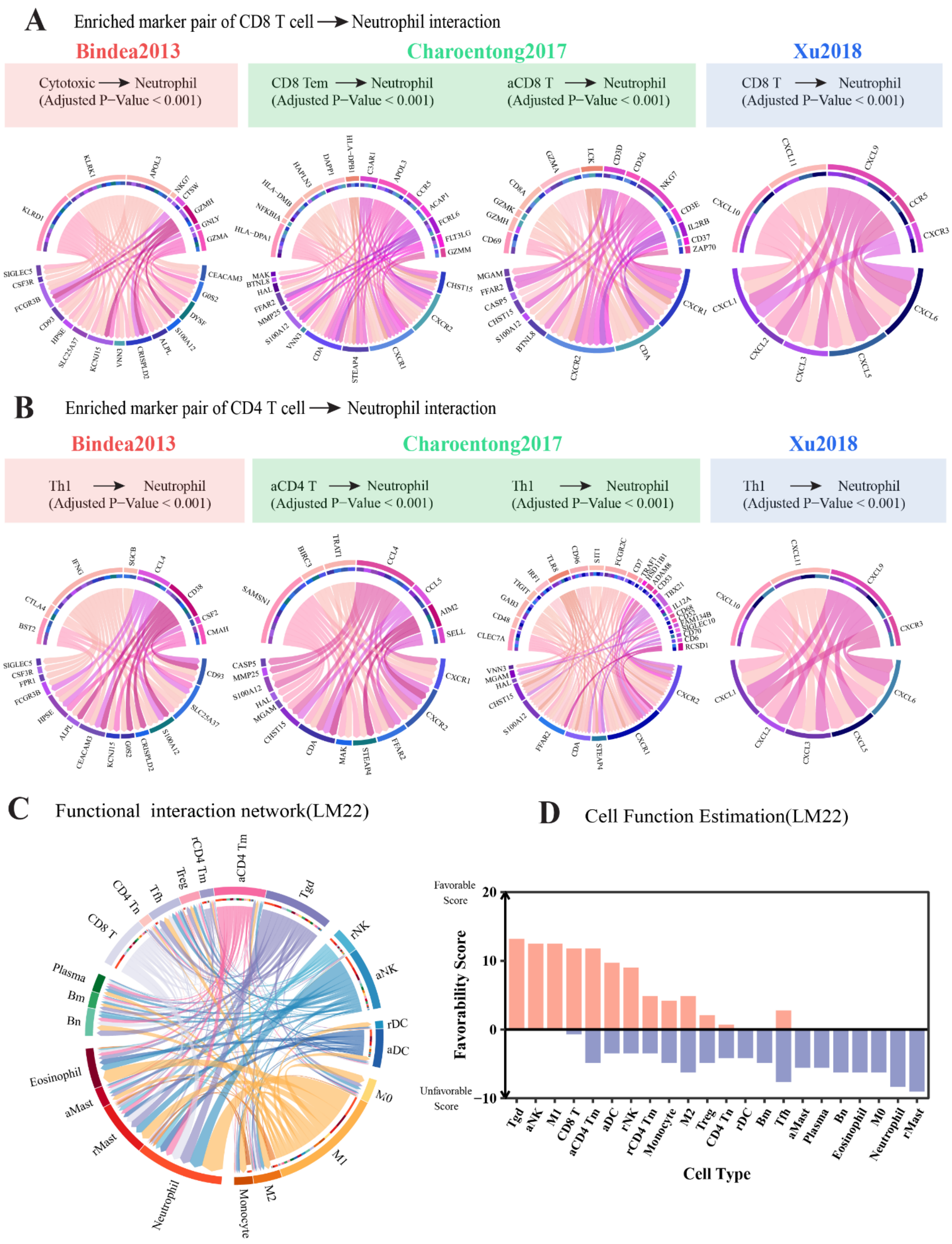
A
Enriched marker pair of CD8 T cell → Neutrophil interaction
Bindea2013
Charoentong2017
Xu2018
Cytotoxic → Neutrophil (Adjusted P−Value < 0.001)
CD8 Tem → Neutrophil (Adjusted P−Value < 0.001)
aCD8 T → Neutrophil (Adjusted P−Value < 0.001)
CD8 T → Neutrophil (Adjusted P−Value < 0.001)
B
Enriched marker pair of CD4 T cell → Neutrophil interaction
Bindea2013
Charoentong2017
Xu2018
Th1 → Neutrophil (Adjusted P−Value < 0.001)
aCD4 T → Neutrophil (Adjusted P−Value < 0.001)
Th1 → Neutrophil (Adjusted P−Value < 0.001)
Th1 → Neutrophil (Adjusted P−Value < 0.001)
C
Functional interaction network(LM22)
CD8 T
CD4 Tn
Tfh
Treg
rCD4 Tm
aCD4 Tm
Tgd
rNK
aNK
rDC
aDC
M0
M1
M2
Monocyte
Neutrophil
rMast
aMast
Eosinophil
Bn
Bm
Plasma
D
Cell Function Estimation(LM22)
Favorable Score
Unfavorable Score
Favorability Score
20
10
0
−10
Tgd
aNK
M1
CD8 T
aCD4 Tm
aDC
rNK
rCD4 Tm
Monocyte
M2
Treg
CD4 Tn
rDC
Bm
Tfh
aMast
Plasma
Bn
Eosinophil
M0
Neutrophil
rMast
Cell Type

### Figure S3 Evaluation of TimiGP robustness with distinct cell type annotations, related to Figure 3

(A-B) Analogous functional interactions identified by distinct cell type annotations

Chord diagram of enriched marker pairs represents (A) CD8 T cell → Neutrophil and (B) CD4 T cell → Neutrophil functional interactions. While cell-type signatures are divergent(Bindea2013, red; Charoentong2017, green; Xu2018, blue), similar cell interactions are identified as functional interactions. In the chord diagram, the arrow represents enriched marker pairs of functional interactions from favorable cell-type markers (color of the outer ring and arrow) to unfavorable cell-type markers (color of the inner ring). Significance is calculated by the TimiGP methods based on enrichment analysis.

(C-D) Application of TimiGP to metastatic melanoma based on CIBERSORT LM22 cell type markers

(C) Chord diagram of all functional interactions. The arrow represents functional interactions from favorable cell type(color of the outer ring and arrow) to unfavorable cell type(color of the inner ring). The wider the arrow is, the smaller the adjusted p-value is.

(D) Bar plot of the favorability score to evaluate each cell type's favorable(orange) or unfavorable(blue) role in anti-tumor immunity and prognosis.

All analysis has been performed with the RNA-seq of metastatic melanoma in the Cancer Genome Atlas(TCGA_SKCM06). Abbreviation of cell types: Bn, Naïve B cell; Bm, Memory B cell; Plasma, Plasma cell; T, T cell; CD4 Tn, Naive CD4 T cell; Tfh, Follicular Helper T cell; Treg, Regulatory T cell; rCD4 Tm, Resting Memory CD4 T cells; aCD4 Tm, Activated Memory CD4 T cells; Tgd, Gamma delta T cell(Tγδ); rNK, Resting Natural killer (NK) cell; aNK, Activated NK cell; rDC, Resting dendritic cell (DC); aDC, Activated DC; M0, non-activated(M0) Macrophages; M1, pro-inflammatory(M1) Macrophages; M2, anti-inflammatory(M2) Macrophages; rMast, Resting mast cell; aMast, Activated mast cell.

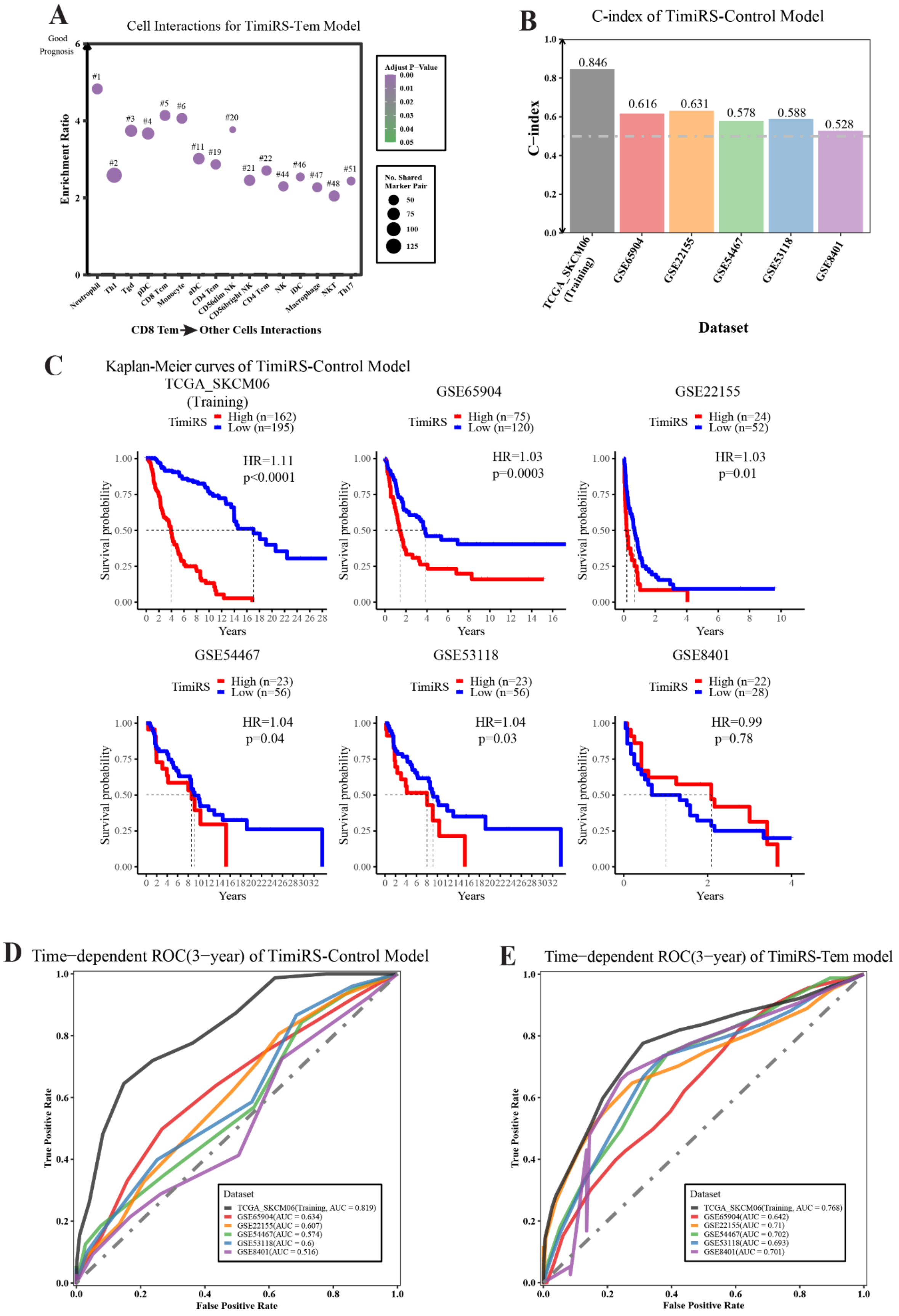
A
Cell Interactions for TimiRS-Tem Model
Good Prognosis
Enrichment Ratio
Adjust P−Value
No. Shared Marker Pair
CD8 Tem ➔ Other Cells Interactions
B
C-index of TimiRS-Control Model
C−index
0.846
0.616
0.631
0.578
0.588
0.528
TCGA_SKCM06 (Training)
GSE65904
GSE22155
GSE54467
GSE53118
GSE8401
Dataset
C
Kaplan-Meier curves of TimiRS-Control Model
TCGA_SKCM06 (Training)
TimiRS High (n=162) Low (n=195)
HR=1.11
p<0.0001
GSE65904
TimiRS High (n=75) Low (n=120)
HR=1.03
p=0.0003
GSE22155
TimiRS High (n=24) Low (n=52)
HR=1.03
p=0.01
GSE54467
TimiRS High (n=23) Low (n=56)
HR=1.04
p=0.04
GSE53118
TimiRS High (n=23) Low (n=56)
HR=1.04
p=0.03
GSE8401
TimiRS High (n=22) Low (n=28)
HR=0.99
p=0.78
Survival probability
Years
D
Time−dependent ROC(3−year) of TimiRS-Control Model
True Positive Rate
False Positive Rate
Dataset
TCGA_SKCM06(Training, AUC = 0.819)
GSE65904(AUC = 0.634)
GSE22155(AUC = 0.607)
GSE54467(AUC = 0.574)
GSE53118(AUC = 0.6)
GSE8401(AUC = 0.516)
E
Time−dependent ROC(3−year) of TimiRS-Tem model
True Positive Rate
False Positive Rate
Dataset
TCGA_SKCM06(Training, AUC = 0.768)
GSE65904(AUC = 0.642)
GSE22155(AUC = 0.71)
GSE54467(AUC = 0.702)
GSE53118(AUC = 0.693)
GSE8401(AUC = 0.701)

**Figure S4 Better performance of the TimiRS-Tem model than TimiRS-Control model in independent validation sets, related to Figure 4**

(A) Dot plot showing the CD8 Tem → Other Cells functional interactions with strong confidence(BH-Adjust.P.Value < 0.0001). These functional interactions were used to construct TimiRS-Tem model. The x-axis shows the other cells unfavorable compared to CD8 Tem. Text labels the rank based on the adjusted P-value in all functional interactions.

The analysis has been performed with the RNA-seq of metastatic melanoma in the Cancer Genome Atlas(TCGA_SKCM06). Abbreviation of cell types in Charoentong2017 annotation: Tem, Effector memory T cell; Th1, Type 1 Helper T cell; Tgd, Gamma delta T cell(Tγδ); pDC, Plasmacytoid dendritic cell; Tcm; Central memory T cells; aDC, Activated dendritic cell; NK, Natural killer cell; iDC, Immature dendritic cell; NKT, Natural killer T cell; Th17, Type 17 Helper T cell.

(B-D) Performance of the TimiRS-Control model in training and independent validation sets

(B) Bar plot of C-index showing the performance of the TimiRS-Control model in training(TCGA_SKCM06) and five independent validation datasets(GSE65904, GSE22155, GSE8401, GSE54467, GSE53118).

(C) Kaplan-Meier curves(right) showing the stratification ability of the TimiRS-Control model. The Hazard Ratio and the corresponding P-value are calculated by the univariate cox regression model that fits the TimiRS as a continuous variable.

(D) Time-dependent ROC(3 years) and AUC showing the evaluation of the TimiRS-Control model.

(E) Time-dependent ROC(3 years) and AUC showing the evaluation of the TimiRS-Tem model.

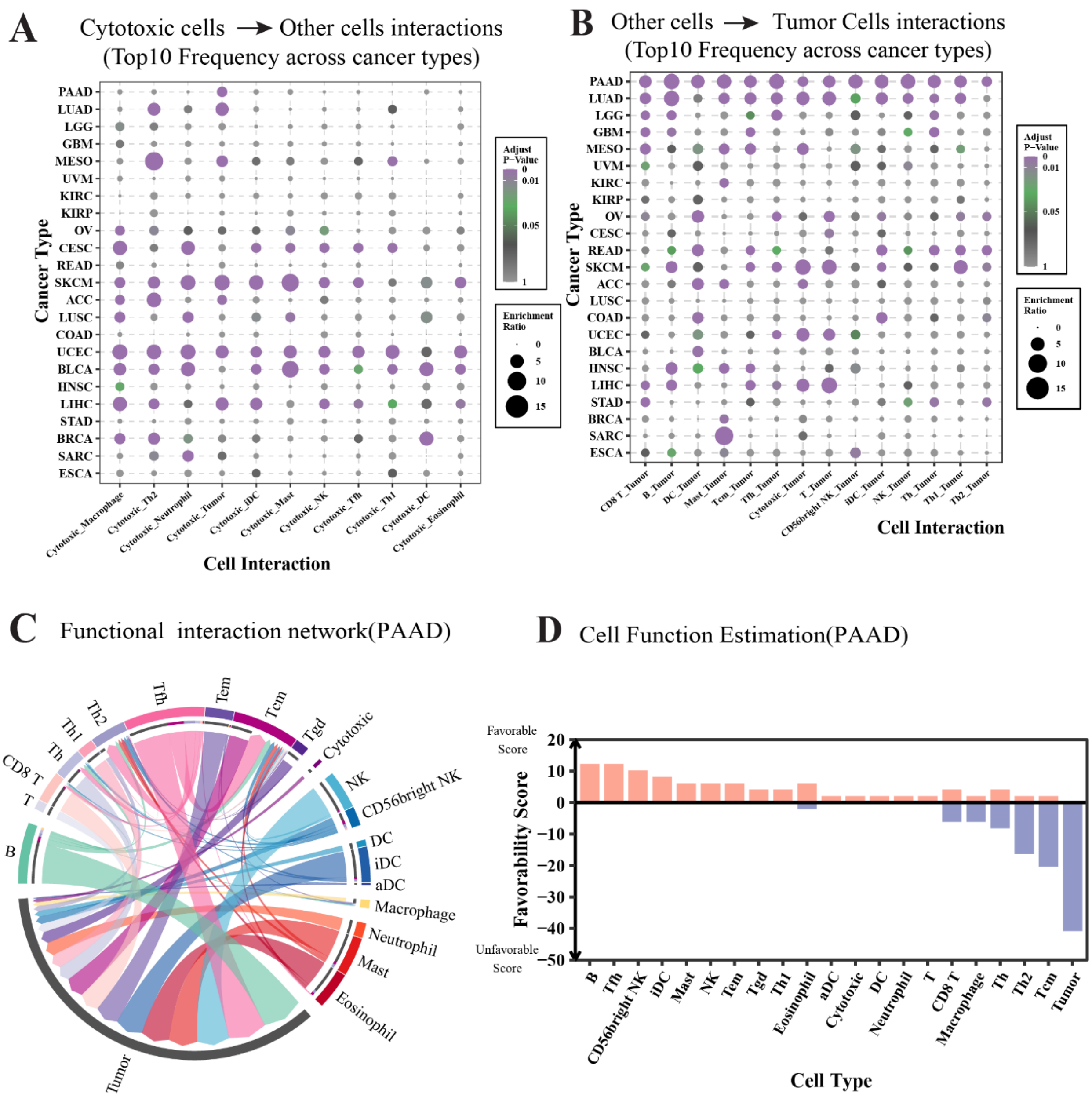


**Figure S5 Pan-cancer functional interactions related to control cells and TimiGP analysis of pancreatic adenocarcinoma, related to Figure 6**

(A-B) Dot plot showing the top 10 functional interactions related to cytotoxic cells(A) or tumor cells(B) in 23 cancer types. A higher ranking is given to the functional interaction involved in cytotoxic cells or tumor cells identified in more cancer types. Significance is BH-adjusted $p < 0.05$.

(B-C) Application of TimiGP to pancreatic adenocarcinoma in the Cancer Genome Atlas(TCGA_PAAD)

(B) Chord diagram of all functional interactions. The arrow represents functional interactions from favorable cell type(color of the outer ring and arrow) to unfavorable cell type(color of the inner ring). The wider the arrow is, the smaller the adjusted p-value is.

(C) Bar plot of the favorability score to evaluate each cell type's favorable(orange) or unfavorable(blue) role in anti-tumor immunity and prognosis.

Abbreviation of cell types in Bindea2013 annotation: B, B cell; T, T cell; Th, Helper T cell; Th1, Type 1 Th; Th2, Type 2 Th; Tfh, Follicular Th; Tem, Effector memory T cell; Tcm, Central memory T cell; Tgd, Gamma delta T cell(Tγδ); Cytotoxic, Cytotoxic cell(common cytotoxic features of anti-tumor CD8 T cells, Tγδ, and NK cells); NK, Natural killer cell; DC, Dendritic cell; iDC, Immature DC; aDC, Activated DC; Mast, Mast cell; Tumor, Tumor cell.

A Rationale: TIME balance affects clinical outcomes

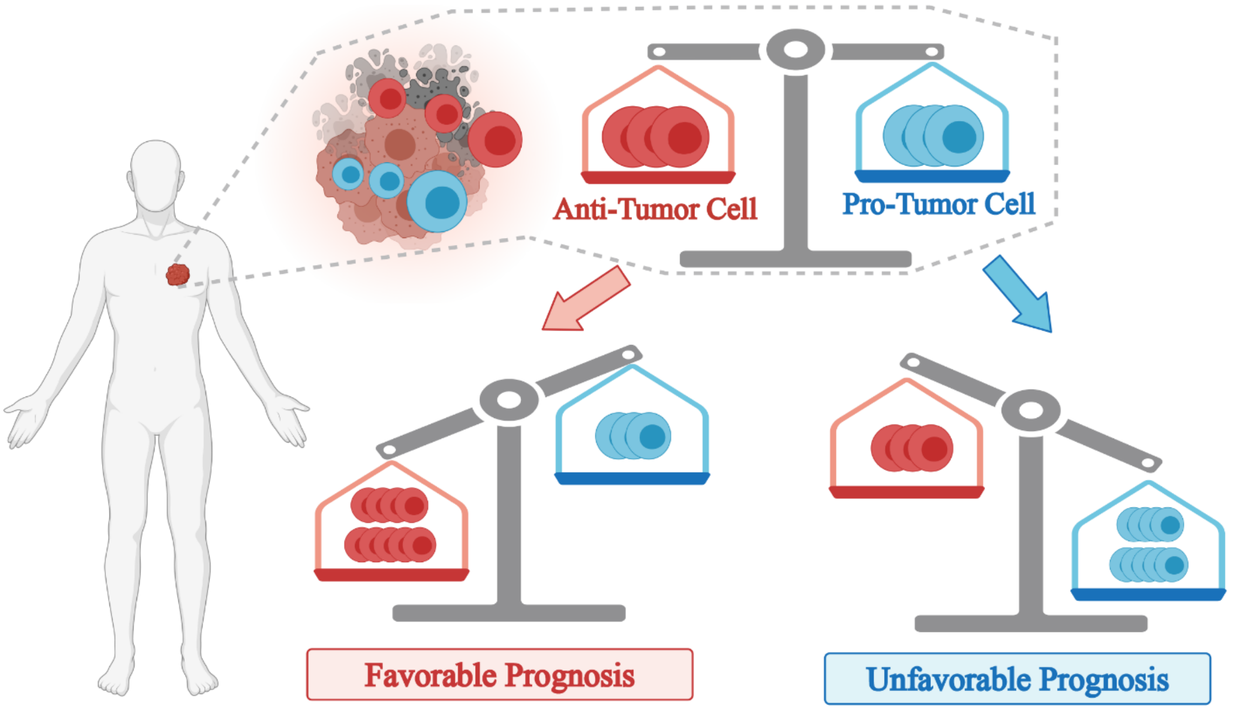


B Summary of TimiGP Applications

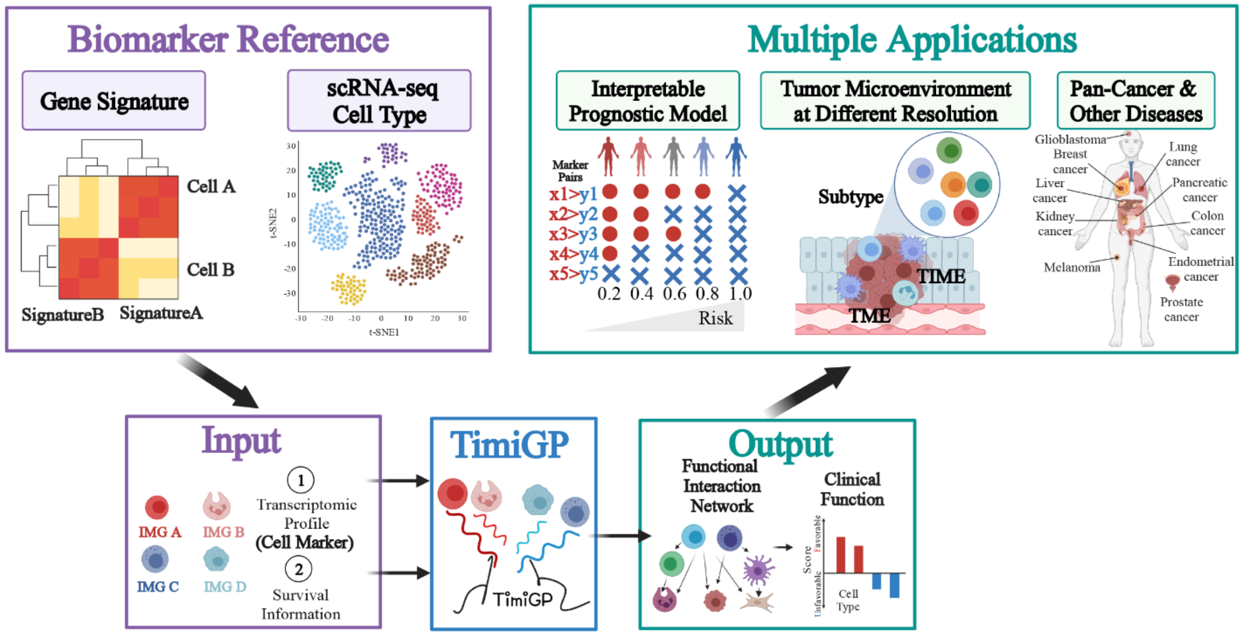


**Figure S6 Summary of TimiGP Rationale and Applications, related to Discussion**
(A) Schema of functional interactions concept based on the immune balance. The tumor immune microenvironment(TIME) is a balance between anti-tumor and pro-tumor immune cells. If the function of anti-tumor cell types is more vital than the pro-tumor cells (e.g., higher abundance, higher marker expression), the TIME is associated with favorable patients' prognosis; otherwise, it is associated with unfavorable patients' prognosis.

(B) Summary of TimiGP Applications. TimiGP is designed to infer the functional interaction network and clinical function of immune cells. Based on the resulting immunological insights, The method is able to facilitate the development of prognostic models. Taking advantage of different biomarker references derived from bulk and single-cell RNA-seq, TimiGP can be applied to investigate the entire tumor microenvironment or cell subpopulations. With transcriptome data in different cancers or diseases, the TimiGP framework is applicable to pan-cancer analysis or other diseases.
